# Local and long-distance organization of prefrontal cortex circuits in the marmoset brain

**DOI:** 10.1101/2021.12.26.474213

**Authors:** Akiya Watakabe, Henrik Skibbe, Ken Nakae, Hiroshi Abe, Noritaka Ichinohe, Muhammad Febrian Rachmadi, Jian Wang, Masafumi Takaji, Hiroaki Mizukami, Alexander Woodward, Rui Gong, Junichi Hata, David C. Van Essen, Hideyuki Okano, Shin Ishii, Tetsuo Yamamori

**Author notes:** Correspondence. Email: Akiya Watakabe, Henrik Skibbe, Tetsuo Yamamori.

## Abstract

Prefrontal cortex (PFC) has dramatically expanded in primates, but its organization and interactions with other brain regions are only partially understood. We performed high-resolution connectomic mapping of marmoset PFC and found two contrasting corticocortical and corticostriatal projection patterns: “patchy” projections that formed many columns of submillimeter scale in nearby and distant regions and “diffuse” projections that spread widely across the cortex and striatum. Parcellation-free analyses revealed representations of PFC gradients in these projections’ local and global distribution patterns. We also demonstrated column-scale precision of reciprocal cortico-cortical connectivity, suggesting that PFC contains a mosaic of discrete columns. Diffuse projections showed considerable diversity in the laminar patterns of axonal spread. In mice, columnar projections were much less conspicuous, underscoring the importance of the primate model. Altogether, these fine-grained analyses reveal important principles of local and long-distance PFC circuits in marmosets and provide insights into the functional organization of the primate brain.

## Introduction

Prefrontal cortex (PFC) plays a central role in orchestrating the activities of various brain regions to achieve complex behaviors involving cognitive and emotional functions, and its malfunction contributes to diverse mental disorders^1,2^. PFC contains a mosaic of distinct cortical areas, with an estimated 45 PFC areas out of 180 total neocortical areas in humans and 35 PFC areas out of 130 total in macaques^3^; the marmoset has 26 PFC areas out of a 117-area parcellation^4^. PFC has a complex pattern of connections with various cortical and subcortical regions^1,2,5,6^. However, our knowledge of PFC connectivity remains fragmentary, particularly for quantitative connectivity data. An existing retrograde tracer database covers most but not all PFC subregions^5,7^, and no quantitative analysis has been reported using anterograde tracers in PFC.

High resolution quantitative anterograde tracer data is especially desirable for primate neocortex, where columnar and modular (periodic) organization have been studied extensively, particularly in visual cortex. Columnar organization generally reflects commonalities along the radial axis (from white matter to pia), exemplified by orientation columns and ocular dominance columns in area V1 and tuning for other features in extrastriate visual areas^8–10^. Modular organization is often associated with patchy anatomical connections in the tangential domain that correlate with repeating representations of a columnar feature such as a particular orientation or eye dominance^8,9^. However, because preferred orientation is represented as a smoothly changing variable, a single ‘column’ in visual cortex is more a conceptual abstraction than a discrete 3D cortical domain whose full extent can be precisely demarcated. In contrast, striking examples suggestive of a different type of columnar organization have been reported using anterograde tracer injections in PFC and other association cortex regions, but major questions remain as to whether such patterns reflect discrete, segregated modules vs highly overlapping connectivity profiles, whether they are predominantly patchy vs stripe-like, and whether their origins and terminations are consistently columnar or are often layer-specific^11–14^.

In the present study, we provide evidence on these and other issues using a dataset based on 44 anterograde and 13 retrograde tracer injections into PFC and adjacent frontal lobe regions in common marmosets. The marmoset is an increasingly popular non-human primate model for neuroscience studies^15–22^. Its cortex is one-tenth the size and far less convoluted than macaque cortex, yet it contains the frontal eye field (FEF), V5 (MT), and granular PFC, all common to primates but lacking clearly defined homologues in rodents^1,20,21^. The high volumetric resolution and sensitive axonal detection of our approach allowed us to systematically analyze patchy (and columnar) corticocortical and corticostriatal axonal projections plus a complementary pattern of diffuse projections generally restricted to one or a few layers. By combining anterograde and retrograde double-tracing in some animals, we demonstrated a striking reciprocity of patchy cortical connectivity. We also mapped the topographic organization of PFC connectivity to multiple regions of association cortex, in a pattern similar but not identical to that revealed by stimulation-fMRI mapping of macaque lateral PFC^23^. Our datasets are freely accessible (https://dataportal.brainminds.jp/; see also^24^) and add to a growing collection of publicly available marmoset neuroscience-related datasets^25^.

## Results

### Serial two-photon tomography imaging in the marmoset brain

A key to our project was the implementation of serial two-photon tomography^26–28^ in the marmoset brain. Combined with the Tet-enhanced adeno-associated virus (AAV) vector system, we achieved efficient detection of even sparsely distributed axon fibers across the entire brain. The high quality of our volume imaging is illustrated in the reconstruction of columnar axonal spread in the cortex for an exemplar dorsolateral PFC (dlPFC) injection (Fig. 1A-1D and Supplementary Video 1 and Supplementary Video 2). There were many conspicuous sites of dense axonal convergence (e.g., Figs 1A-1D, red arrowheads and white arrows) adjacent to regions with only sparse axonal signals (Fig. 1B, right panels, compare “in” vs “out”). Depending on section obliquity relative to the radial axis, a single section may capture most of a projection column (Fig. 1B) or only part of it (Fig. 1A), but in all cases serial sections revealed the full 3D pattern (Fig. 1C, D). Registration fidelity to the common template space^24^ was estimated to be 100-200 μm (Fig. S1A; see STAR Methods), which enabled reliable automatic anatomical annotations (Fig. S1B) and integration across multiple datasets. The slice interval was set at 50 μm, half that used for the anterograde data from mice^27^, thereby enabling smooth conversion of cortical layers to a stack of flatmaps (“flatmap stack”) (Fig. S1C) even in regions of cortical curvature (e.g., frontal pole, dorsomedial convexity) (see Fig. S1E for distortion of intracortical areas). We separated the axon signals from background noise (e.g., lipofuscin granules; Fig. S1F) using a machine learning-based algorithm, achieving five orders of magnitude range of signal intensity. Pseudocolor scaling of logarithmic values revealed both convergence of axons into patches and diffusely spread weak signals (Fig. 1E-1G). The columnar nature of these exemplar patches is particularly evident in an oblique 3D view of PFC (Fig. 1G). Importantly, the patches were not randomly scattered in the tangential domain, but are mostly grouped into rows, consistent with a stripe-like arrangement^12^ if the data were analyzed at lower resolution (see also Fig. 2A).

**Figure 1:**
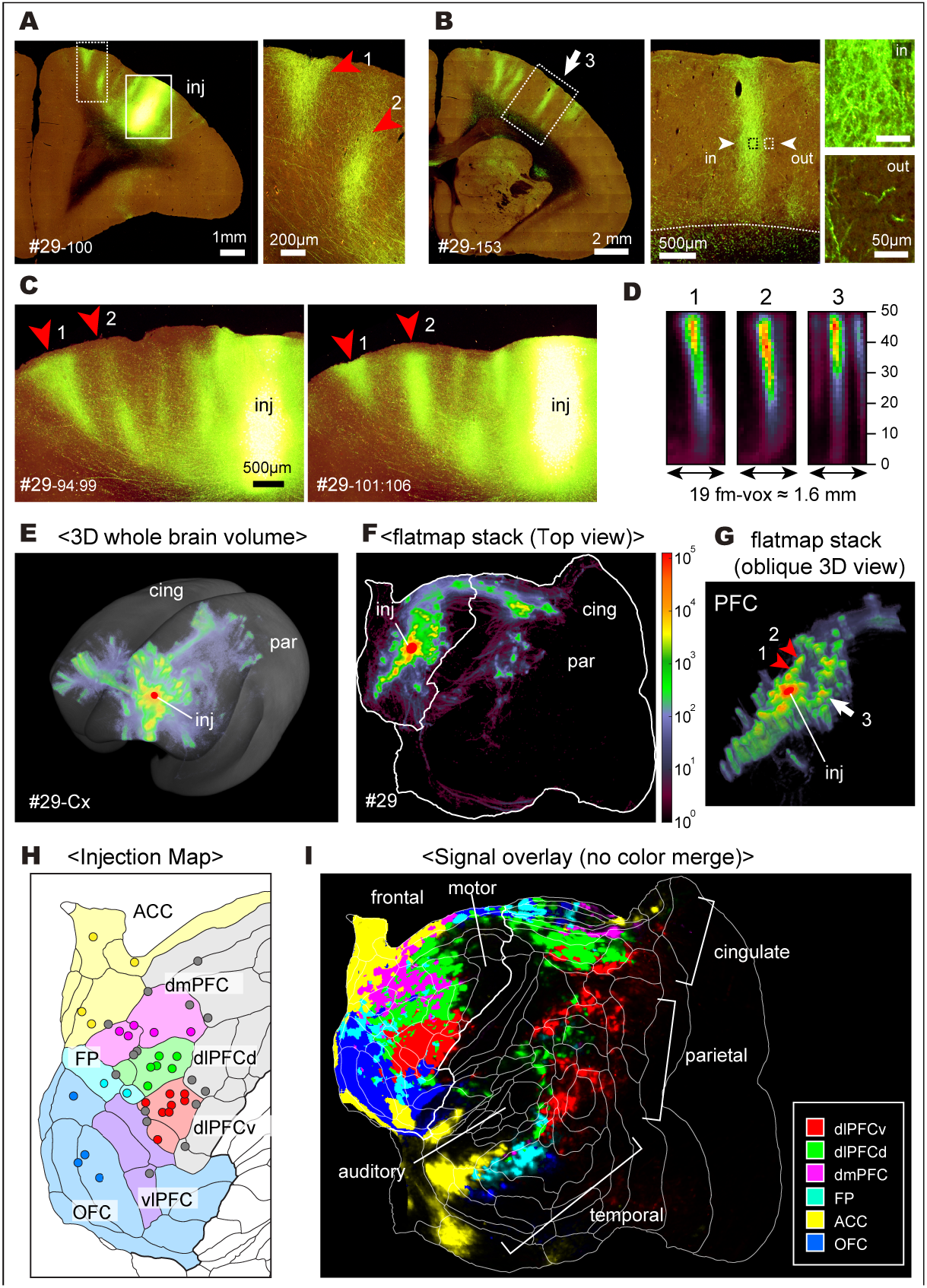
Patchy/columnar projections from marmoset PFC imaged in 3D. **A**, Original section image (#29-100 indicates section 100 of case #29) around the injection site (inj), cut slightly oblique to the radial axis. Note that we could identify the AAV-transduced cell bodies in the unsaturated channel (Fig. S1D). Two red arrowheads (1 and 2) correspond to those in panels C and G. **B**, Another example of a columnar axonal convergence (white arrow 3) shown at three different magnifications of a section cut mostly parallel to the radial axis. **C**, Maximum intensity projection (MIP) views for six serial slices showing radial extensions of columns 1 and 2, before and after section 100. **D**, Three-dimensional (3D) reconstruction of columns 1, 2 and 3 in the flatmap stack shown as a MIP side view. This and other similar images (e.g., panels E, F and G) use the positive half of the ‘Videen’ palette for pseudocolor intensity scaling (see STAR Methods). fm-vox; flatmap-voxel (see STAR Methods). **E**, 3D reconstruction of the cortical tracer signals in the STPT template space. The log value and pseudocolor intensity scaling are used to show both strong columnar protrusions and diffuse spread over broad regions of the cortex. The injection site (inj) is shown by a red circle. cing; cingulate field. par; parietal field. **F**, An MIP top view of the flatmap stack image for the left cortex for the data shown in panel E. **G**, A 3D view of the flatmap stack around the injection site. Example columnar projections (1-3) are indicated by the arrow and arrowheads. **H**, Summary figure of the injections analyzed in this study (also see Fig. S1G). Injections centered well inside a core domain (darker colors) were used for analyses of regional differences in connectivity. Injection sites shown in gray were excluded from this regional analysis either because they position at the boundary or in the premotor (PM) areas, but were included in other parcellation-free analyses. **I**, Overlay image of projections showing a systematic distribution pattern (linear scale and no color merge; see also Fig. S1H and I). Subregion abbreviations: dlPFCd (dorsolateral PFC-dorsal), green; dlPFCv (dorsolateral PFC-ventral), red; dmPFC (dorsomedial PFC), magenta; FP (frontopolar cortex), cyan; ACC (anterior cingular cortex), yellow; OFC (orbitofrontal cortex), blue.

**Figure 2:**
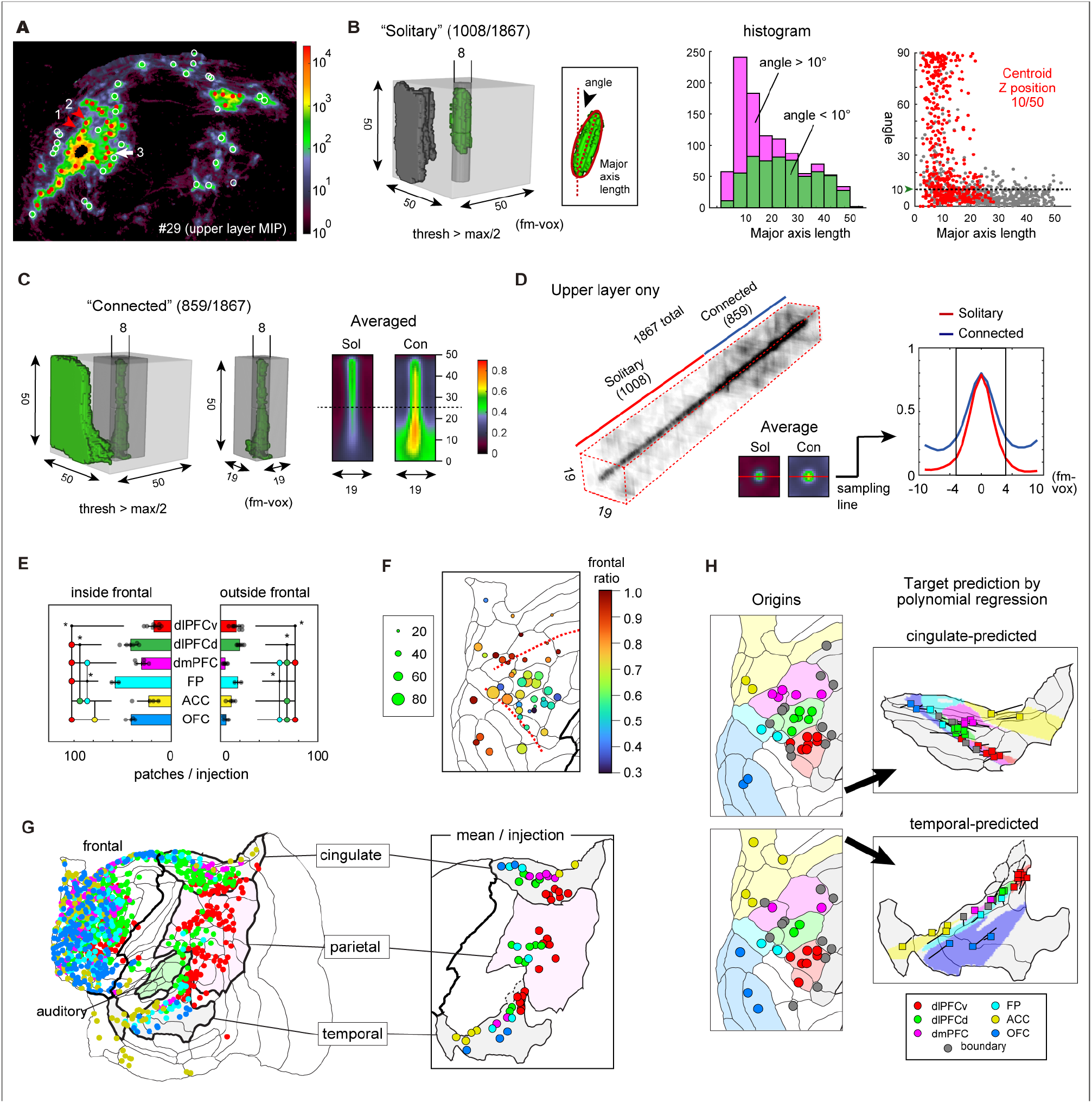
Topographic distribution of columnar cortical projections from PFC subregions. **A**, Detection of “columnar patches” in the cortical flatmap stack as the local maxima. The red dots and white circles indicate patches with peak values that are >1/10 (10^3^) and >1/100 (10^2^) of the standardization value (10^4^), respectively. **B**, 3D reconstruction of binarized tracer signals (threshold at half the maximum value of the upper cylinder). We identified patches as connected or solitary based on whether the clump was (panel C) or was not (panel B) connected to surrounding labeled regions (see STAR Methods for details). The clumps for solitary patches were approximated by an ellipsoid, and its major axis length and the angle of deviation were measured. The histogram shows the distribution of major axis lengths of those with small (<10°; green) and large (>10°; magenta) angles. The red dots in the scatter plot show the clumps centered in the uppermost (10/50) compartment. fm-vox; flatmap-voxel (see STAR Methods). **C**, an example of the “connected” patches. The averaged laminar profiles for the solitary (Sol) and the connected (Con) patches show the averages of MIP view of the central 4 (thickness) x19 (width) x50 (depth) voxel boxes. **D**, The non-binarized original tracer signals in the upper layer compartments were averaged to show their 19 × 19-fm-vox planar spreads for 1867 columnar patches, which were aligned in 3D to show the central convergence. These planar images were further averaged for the two groups (Sol and Con) and the intensities were measured along the red sampling lines. **E**, The average number of columnar patches per injection for six PFC subregions. The colored circles indicate statistically significant differences (ANOVA with Tukey’s honest significant difference test, p<0.05) between the subregions of these colors and the subregions left (or right) of the circles. **F**, The number of columnar patches (in circle diameter) and the ratio of the frontal /total patch numbers (shown by color scale) for each injection are displayed. **G**, Distribution of the columnar patches for the six PFC subregions in four association area fields and early auditory areas (see also Fig. S3D). The averaged positions of the columnar patches in the cingulate, parietal and temporal fields are shown in the right panel. **H**, Visualization of topographic relationships in the cingulate and temporal fields. The line bars indicate the deviations of the true positions from the predicted ones. The injections shown on the left had at least one columnar patch in the projection field.

Fig. 1H and Fig. S1G show the estimated locations of injection sites on the cortical flatmap that includes a 117-area cortical parcellation (Fig. S2) based on an architectonic analysis of a single hemisphere^4,22^. The areal boundaries provide a useful reference frame, but our regional analysis was based on clustering the atlas areas into 6 larger PFC ‘core’ subregions identified by geographic location, shaded in different pastel colors in Fig. 1H and assigned geographic labels as indicated in the figure and legend. Fig. 1I represents the overlay of projections from the six core PFC subregions using linear scaling. The projections from different subregions were largely segregated in the three posterior association area fields and had very different weightings, with dlPFCd projections (green) dominating the cingulate field, dlPFCv projections (red) dominating the parietal field, and OFC and ACC projections dominating the temporal field. There is a quasi-orderly topographic representation in each of the association fields (see below and Fig. S1H and S1I). Using this coverage, the distribution patterns and principles of patchy/columnar projections and diffuse spread are examined in detail below.

### Columnar projections recapitulate PFC topographic gradients locally in each association field

To characterize axonal convergence patterns in the cortex in greater detail, we searched for local maxima in the flatmap (Fig. 2A) and identified 1867 patches of strong labeling from all injections combined. To assess how consistently these patchy patterns are aligned as columns parallel to the radial axis, we binarized the tracer signals and determined that more than half (54%, 1008/1867) of the binarized 3D clumps were solitary (disconnected from surrounding label) and reasonably well approximated by ellipsoids. Of the 1008 solitary patches, 569 (56 %) had ellipsoids whose major axes were within 10 degrees of the estimated radial axis and had lengths that varied widely (Fig. 2B, green histogram); we consider all of these to be oriented radially. In the remaining 44 %, most patches were insufficiently elongated to identify a clear axis of orientation; most of them represent isolated patches in superficial layers (Fig. 2B, red dots in scatter plot on right). For the 859 “connected” cases (46 %), the averaged data suggested that most of them were associated with radially oriented signal convergence, especially in the upper layers (Fig. 2C). Indeed, a translucent display of upper-layer portions of all patches aligned as a single stack (Fig. 2D) indicated that patches were consistently narrow in diameter (∼8-10 voxels) for both solitary (“Sol”) and connected (“Con”) cases. This corresponds to ∼672∼840 μm diameter in the flatmap stack (see Fig. S1E). We conclude that the submillimeter connectivity patches, both local and long-distance (including those projecting outside PFC), predominantly have a radially oriented columnar architecture, although intensity profiles can vary considerably across layers, and many patches are restricted to superficial cortical layers. Accordingly, we refer to these as ‘columnar patches’ or ‘columns’.

Each injection generated a variable number of columnar patches within and outside frontal cortex (Fig. 2E, one-way ANOVA, p<0.05). Within frontal cortex, subregions FP, dlPFCd, and OFC injections averaged ∼40 - 60 patches/injection, whereas dlPFCv, dmPFC, and ACC had significantly fewer. Outside frontal cortex, dlPFCd, dlPFCv, and FP averaged ∼10-20 patches/injection and the others had fewer still. A map of the total number of columnar patches (circle diameter) and the frontal/total ratio (circle hue) for each injection locus showed total numbers highest for dorsolateral injections and high frontal ratios mainly in orbitomedial subregions (Fig. 2F), suggesting regional differences in the number and distribution of columnar projections.

A more detailed examination revealed a finer-grained pattern for projections from PFC to columnar patches in three association fields in cingulate, parietal and temporal cortex (Fig. 2G). In both cingulate and temporal cortex, we observed partially segregated columnar patches from all six subregions, suggesting that PFC topography may be recapitulated in each of these fields. Indeed, we could infer the approximate target coordinates based on the injection coordinates using polynomial regression models (Fig. S3A and 3B), which revealed a skewed transformation of the PFC topographic gradients (Fig. 2H). The projections to parietal cortex differed in that columnar patches arose only from dorsolateral injections (dlPFCd, dlPFCv, FP-red, green, cyan, Fig. 2G). A seed analysis using all injections showed segregated projections from the anterior and posterior compartments of the dorsolateral frontal lobe (Fig. S3C). We also found columnar patches in early auditory areas (the core and belt) originating from two portions of dlPFCd and dlPFCv (Fig. S3D). We propose that the patchy columnar projections from the PFC take parallel pathways toward these extra-frontal fields, where they recapitulate key aspects of PFC topography in region-specific ways. Fig. S3E and 3F show that these topographic projections are impressively similar to those reported for macaque lateral PFC^23^. We also found orderly relationships in the distribution of columnar patches within frontal cortex (Fig. S3G-I), suggesting extensive mutual connectivity by columnar projections within frontal cortex, with an overall extent that varies systematically across subregions.

### Columnar PFC projections are far less conspicuous in mice than in marmosets

Columnar projections have been considered rare in rodents^29^. To systematically compare them with marmoset, we converted the 3D data of the mouse projectome generated by the Allen Institute for Brain Science using anterograde tracers^26,27^ to the flatmap format and performed the same patch detection analysis as was done for the marmoset. An exemplar mouse OFC injection (Fig. 3A, left, in area ORBm) showed predominantly diffuse projections in the cortex (Fig. 3B). Our patch detection algorithm identified some local maxima (Fig. 3D; right panel in Fig. 3A) but many fewer than from an OFC injection in the marmoset (Fig. 3E and 3F). We performed columnar patch detection for all available mouse datasets with injections into putative prefrontal areas of wild-type mice (Supplementary Table 3). A comparison with the marmoset data (Fig. 3G) suggests that columnar patches are far less conspicuous for inter-areal mouse PFC projections and that the strong within-area patchiness observed for marmosets (e.g., robust columnar projections near the injection site in Fig. 3E and 3F) occurs rarely in mice.

**Figure 3:**
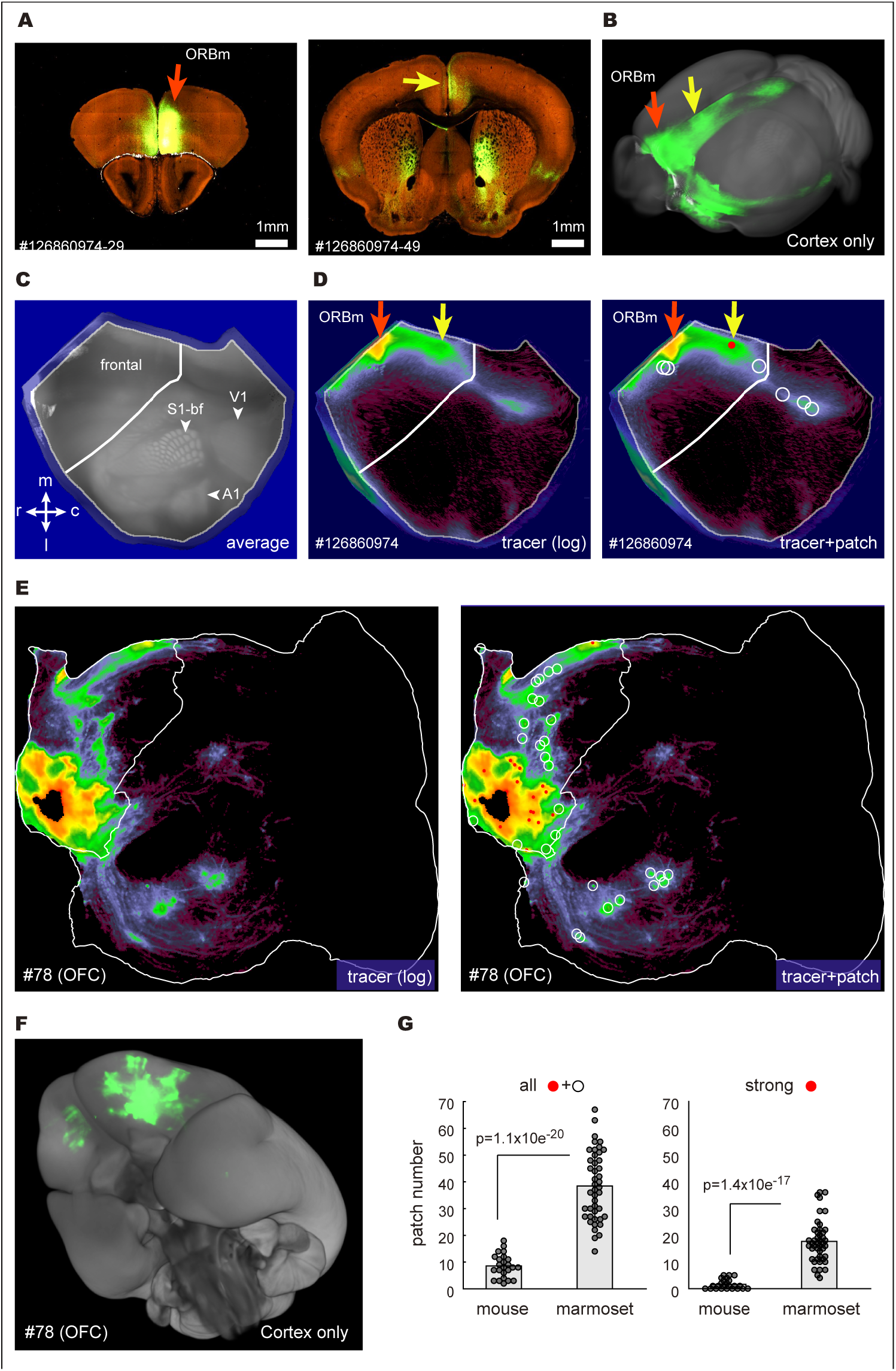
Comparison of mouse and marmoset data for columnar projections. **A**, Left: A coronal section centered on an injection site (red arrow) in ORBm (126860974) retrieved from the Mouse Brain Connectivity Atlas^26,27^. Right: coronal section through a local maximum detected as in panel D (shown by a yellow arrow). **B**, Translucent 3D reconstructed view of the cortical projection data. **C**, Flatmap image for the averaged background fluorescence for cortical landmarks. Owing to spatial distortions, the fringes of the flatmap were excluded from patch detection. The middle white line indicates the border between the motor and somatosensory areas. S1-bf; S1 barrel field. m; medial, l; lateral, r; rostral, c; caudal. **D**, The flatmap image of the ORBm injection in logscale was pseudocolored to show both strong and weak signals. Red dots and white circles on the right indicate columnar patches detected with our algorithm with peak values that are >1/10 and >1/100 respectively of the standardization value (see STAR Methods for details). **E**, Detection of columnar patches for marmoset injection #78 (OFC) using the mean of all layers for comparison with the mouse data. The white line in the middle indicates the border between motor and somatosensory areas. **F**, Translucent 3D reconstructed image of the cortical projection data for case #78 viewed from the bottom. **G**, Comparison of detected patch numbers per injection between the mouse and the marmoset. The right panel compares only the strong signals (shown as red dots). See Supplementary Table 3 for information on the mouse data (24 samples) used in this analysis. All 44 samples were used for the marmoset data.

### Diffuse projections recapitulate PFC topographic gradients globally

The highly distributed pattern of weak anterograde labeling visible using log scaling and gray-scale encoding is shown in Fig. 4A for injections in dlPFCv (top) and dmPFC (bottom) and for 5 additional injections in Fig. S4A. Diffuse projections were even observed in regions lacking any columnar patches (Fig. 4A, green arrows), suggesting that their specificity should be considered separately from that of the columnar patches. Because the diffuse patterns appeared to overlap substantially even across injections into different subregions, we performed a nonnegative matrix factorization (NMF) analysis to identify common components contributing to various patterns. Figs 4B and 4C show the results of NMF, which generated four components (basis images) (NMF_W1, 2, 3 and 4) and associated coefficients (coeff. 1, 2, 3 and 4), which could efficiently and reasonably accurately reconstitute the original images (Fig. S4B). The distinctive features of these components became more apparent when they were paired for comparison (Fig. 4B; separated by a vertical blue bar). One pair (NMF_W1 and NMF_W4) represents mainly the dorsal versus ventral spread, respectively, reminiscent of the dual origin concept proposed by Pandya and coworkers^6^. The second pair (NMF_W2, and NMF_W3) represents the rostral (peripheral) and the caudal spread, respectively, and resembles the antagonism of the “apex transmodal network” versus the “canonical sensory-motor network” (primarily visuo-motor and distinct from the somato-motor network) previously described in the marmoset^30^ based on retrograde tracer data^5^. The subregion-specific patterns are reflected in the coefficient values for these components, namely, coeff. 1 through 4 (Fig. 4C, One-way ANOVA, posthoc Tukey test, *; p<0.05). Notably, dlPFCv was the only subregion with high coeff. 3 values, indicating that NMF_W3 was nearly unique to dlPFCv. In other cases, 2, 3, or 4 subregions had substantial coefficients, suggesting that the positional gradients of injections strongly influence these coefficients.

**Figure 4:**
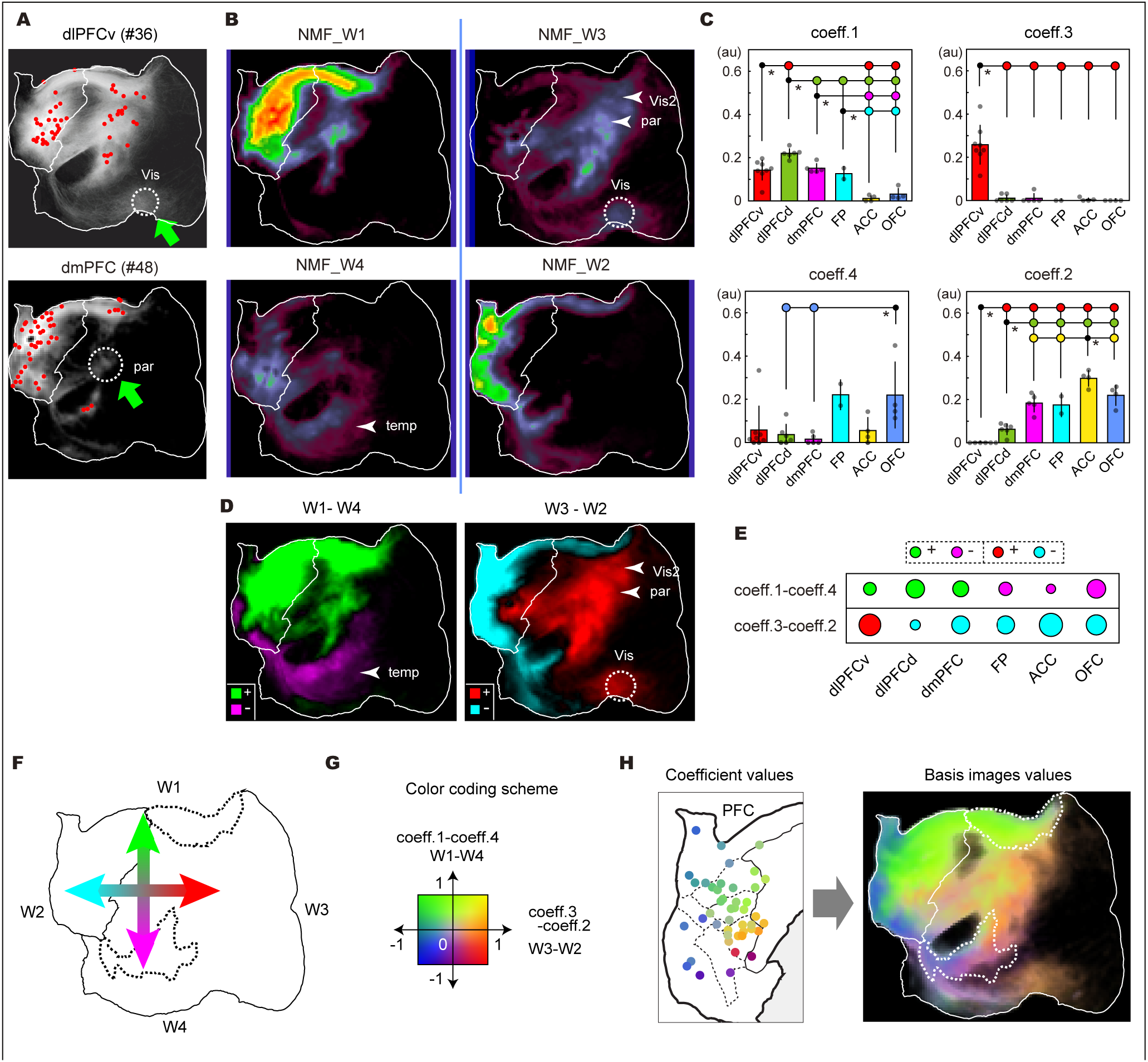
Characterization of diffuse corticocortical projections in log-scale by nonnegative matrix factorization (NMF) analysis. **A**, Log-transformed tracer images for two examples (see Fig. S4A for additional examples). The red dots indicate the detected columnar patches in the same samples. Note the presence of weak signals in the dotted circles indicated by green arrows, where we detected no columnar patches. par, parietal area; Vis, visual area. **B**, Four components (basis images) obtained by the NMF analyses. The intensity is represented in pseudocolor (Videen palette). **C**, Coefficients of each injection, grouped by PFC subregions. Error bars indicate s.d. See the legend in Fig. 2E for statistical significance. **D**, The subtracted values for two pairs of NMF components (NMF_W1-W4 and NMF_W3-W2) are shown with different colors for the positive and negative values (see STAR Methods). **E**, Representation of the subtracted values of the coefficients. The averaged values after subtraction are shown as circle areas with different colors for the positive and negative values. **F**, Schematic representation of the orthogonal gradients observed in panel D. **G**, 2D color map used to represent the variable combinations of the subtracted values for the components and coefficients. **H**, Display of the ratio of the components at different cortical locations (right panel) and coefficients for each injection (left panel) using the 2D color map in panel G. These colors were assigned to regions above the threshold value. Below-threshold regions with almost no axon signals (e.g., early somatomotor, insular and visual areas) are shown black. Note that these colors are not the same as shown in panels D or E.

To visualize the differential contributions of the four parameters in a simpler form, we combined the antagonistic pairs by subtraction, retaining most of the original information in the positive and negative domains (Fig. 4D). Similarly, we combined the coefficients by subtraction (Fig. 4E). This data conversion highlighted the orthogonal relationships of NMF_W1 through W4 (Fig. 4F). Using a 2D color index strategy^23^ to represent the various contributions of W1 through W4 (Fig. 4G) enabled representation of all four components in a single map (Fig. 4H, right panel). A similar strategy revealed the coefficient values associated with each injection point (Fig. 4H, left panel). The gradual progression of hues suggests graded changes in the combination of coefficient values according to injection location, which was modeled by polynomial regression (Fig. S4C, right panel). Since the projection patterns reflect the coefficient values, we infer that the source region mainly projected to target regions similar in hues (compare the left and right panels of Fig. 4H). Note that global gradients and local gradients are not mutually incompatible. As shown in Fig. S4D, injections into three separate locations showed marked differences in the numbers and intensity of patches in cingulate and temporal fields, while maintaining similar topographic relationships. We suggest that PFC gradients are mapped globally across the cortical hemisphere to determine the overall projection patterns but are also mapped locally in each association field to determine the positioning of columnar projections.

### Columnar and diffuse projections have different laminar profiles

The laminar profile of axon terminals has been used to infer hierarchical relationships between connected areas^31–34^. To investigate this issue in our data, we classified the laminar patterns of columnar patches by hierarchical clustering (Fig. 5A and Fig. S5A). Based on the stereotypical patterns reported previously, one might suspect that clusters targeting upper and deeper layers (grouped as UL and DL, respectively) represent feedback connections, whereas those that target widely across the middle layers (ML) might be feedforward or lateral (horizontal) connections. We obtained eight clusters that were combined into DL, ML, and UL groups, with a majority being ML type (green), and examined their areal distribution patterns (Fig. 5B for #47, and Fig. S5B). These examples showed a tendency for similar laminar types to cluster, albeit with considerable intermingling. Next, we examined the laminar profiles of the overall areal projections, including both the columnar patches and the surrounding diffuse projections. Using the 117-area parcellation (Fig. S2A), we selected 4,000 injection-to-target area pairs out of 44×117 possible combinations (78 %) as putatively positive (see STAR Methods) and classified them by hierarchical clustering into four types: UL, ML, DL, and DL2 (Fig. 5C). The incidence of ML laminar types was much lower than for columnar patches in frontal cortex (Fig. 5C, upper panel) and even more so outside frontal cortex (Fig. 5C, lower panel), as is also apparent for the exemplar injection (#47, Fig. 5D). In many areas, the area-wise laminar types differed from the dominant patch types, presumably because strong but restricted columnar signals in the middle layers were dwarfed by diffuse, widespread projections in deep layers (Fig. 5E and 5F). Fig. S5E compares the percentage of the dominant laminar types of the columnar patches with that of the area-wise measurement (left panel), which showed a decrease of the ML type and an increase in DL and DL2 types. Importantly, some areas are reciprocally connected with one another by DL pathways in both directions (Fig. S5F). If these represent genuine inter-areal connections and not just fibers of passage, this pattern would be inconsistent with a traditional hierarchical model in which DL represents a feedback pathway (see Discussion).

**Figure 5:**
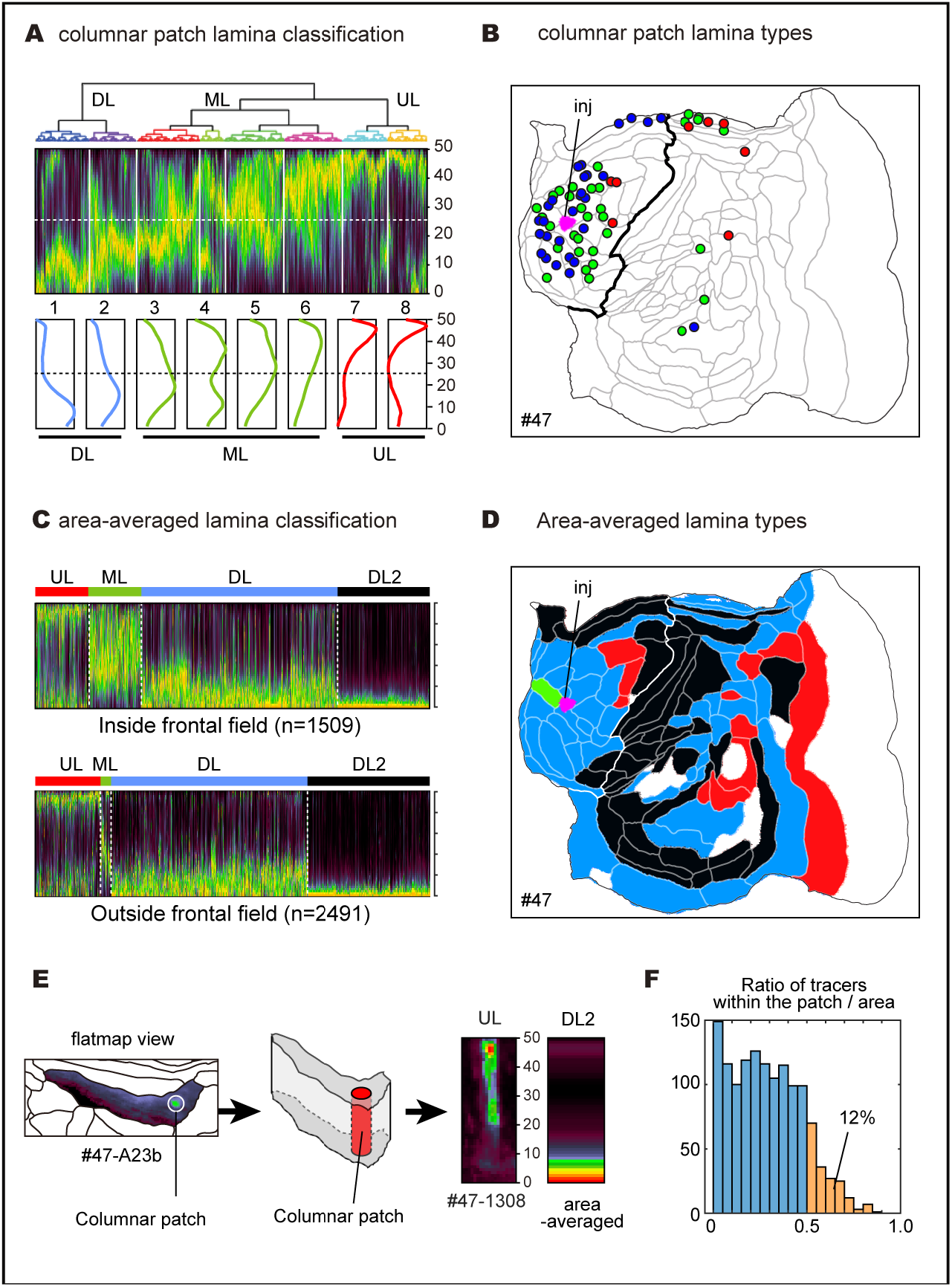
Differential laminar preferences of the columnar patches and diffuse projections. **A**, Classification of the laminar patterns of the columnar patches. Laminar profiles for the 1867 columnar patches were arranged based on the hierarchical clustering. The averaged profiles of eight clusters are shown below, and are further classified into the upper layer (UL), middle layer (ML), and deeper layer (DL) groups. **B**, An exemplar map showing the laminar types (red for UL, green for ML, and blue for DL) of columnar patches for sample #47. The thick line indicates the frontal cortex border. **C**, Classification of the laminar patterns of the area-wise connectivities. Area-based lamina profiling (integrating patchy and diffuse projections within each area) of 4,000 source-target combinations into four groups. **D**, An exemplar map showing the area-averaged laminar types (red for UL, green for ML, blue for DL, and black for DL2) for sample #47. **E**, A schematic diagram explaining the differential lamina patterns for the columnar patches and area averages. An example patch (ID1308 of sample #47) in area A23b is shown. **F**, A histogram showing the ratio of the amounts of tracer within columnar patches vs that for the entire area (diffuse + patchy). Those over 50% (shown in red) constituted only 12% of the 1,325 source-target combinations having columnar patches.

### Reciprocity of corticocortical projections determined by anterograde/retrograde double-tracing

It is well established that most but not all inter-areal corticocortical connections are reciprocal^7,14^, but the degree of spatial precision largely remains to be determined. Thus, we investigated whether both columnar and diffuse projections in the marmoset PFC are associated with reciprocal projections. To test this, we co-injected a non-fluorescent retrograde tracer in 14 cases and compared its distribution pattern with the anterograde tracer by immunostaining. Fig. 6 shows our main findings using case #80 as an exemplar. The retrograde signals exhibited striking colocalization with the anterograde signals by visual inspection (Fig. 6A), in densitometry (Fig. 6B) and cross-correlation analyses (Fig. 6C). Retrogradely labeled neurons occurred in all cortical layers except layer 1 in many patches (Fig. 6A,B), but could be concentrated in deeper layers outside patches (Fig. 6E). Strong columnar patches were consistently associated with moderate to strong retrograde labeling. We also observed a widespread but sparser pattern of retrogradely labeled neurons in locations containing diffuse anterograde label (Fig. 6E). Thus, reciprocal connectivity applies to both patchy columnar and diffuse projections (Fig. 6D).

**Figure 6:**
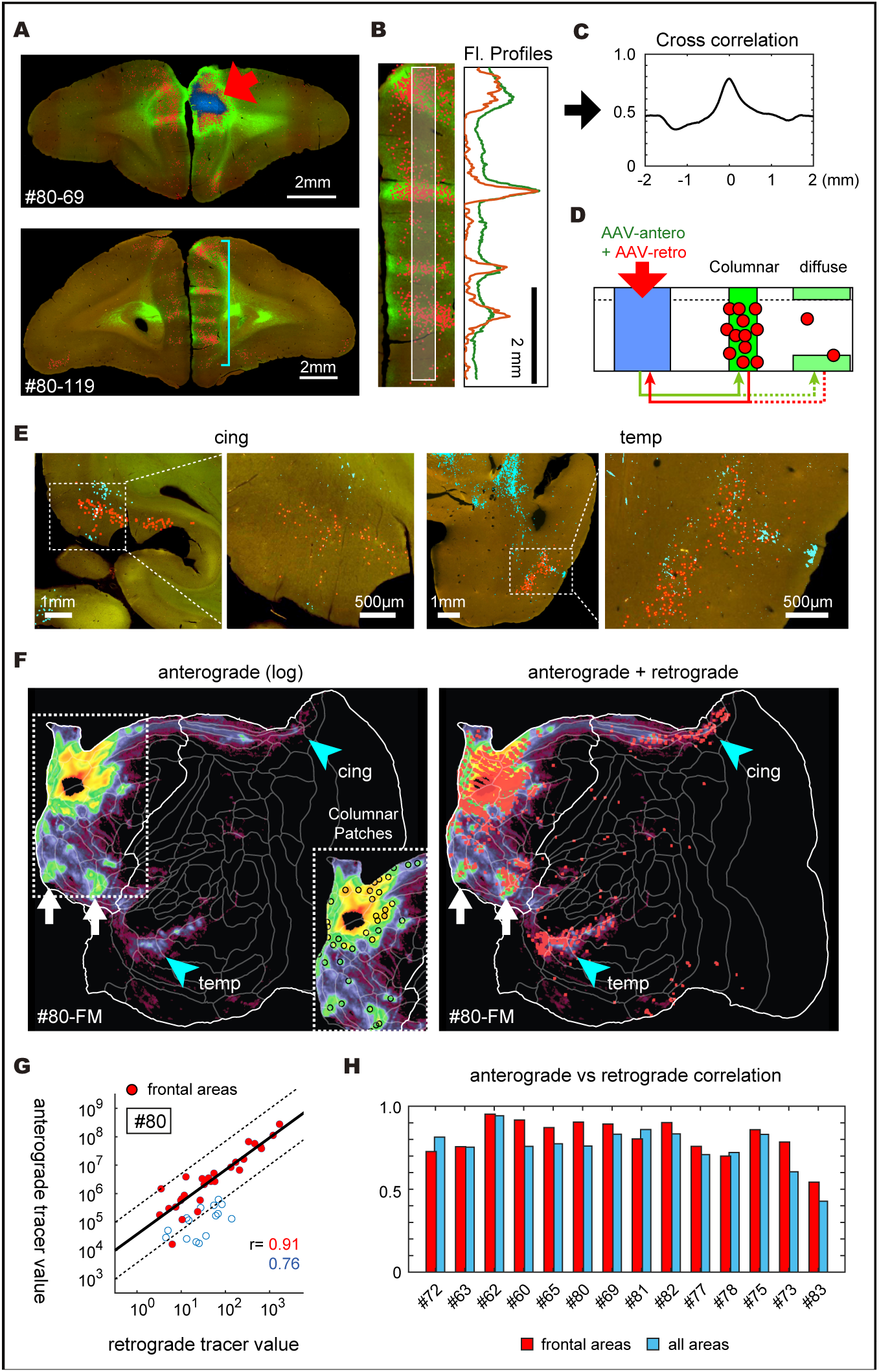
Reciprocal connectivity of corticocortical projections. **A**, Example images showing colocalization of anterograde and retrograde tracers. The injection site is indicated by the blue overlay and a red arrow. Retrogradely-labeled cell nuclei were automatically segmented and overlaid onto the images (red dots). **B**, The fluorescent intensities in the red and green channels were measured along the vertical strip in the middle layers (outlined by white lines) to show colocalization. **C**, Cross-correlation analysis was performed to measure the coincidence of the anterograde and retrograde peaks shown in panel B. **D**, A schematic representation of reciprocal columnar connectivity visualized by co-injection of anterograde and retrograde AAV tracers. **E**, Colocalization of the anterograde (cyan) and retrograde (red) signals in the cingulate and temporal regions. Here, both signals were segmented and contrast-enhanced for better visualization. **F**, Left: Anterograde label alone (pseudocolored in log scale). Inset shows the positions of the columnar patches. Right: retrogradely labeled neurons (red dots) overlaid on the anterograde data. Because we stained only one in ten sections for the retrograde tracers, there were gaps in the flatmap (e.g., around the injection site). **G**, Area-based comparison of anterograde and retrograde signals displayed on a log-log scale. The solid line indicates the best fit for the frontal areas (red dots) and the dashed lines above and below indicate 10-fold differences. The open circles indicate areas outside the frontal field. The correlation coefficients (r) for the frontal areas only and for all areas combined are shown in red and blue lettering, respectively. **H**, Correlation coefficients of the retrograde and anterograde signals in the frontal areas only (red bars) and for all areas (blue bars). Two injections that intruded into white matter were excluded from the analysis. The 5 cases with noisy retrograde signals (#77 - #83 on the right) tended to show lower correlations (see Fig. S6).

To quantify these relationships, we projected the retrograde data into a flatmap for comparison with the anterograde data (Fig. 6F). We confirmed that strong anterograde signals were consistently associated with clusters of retrograde signals (e.g., Fig. 6F, white arrows, compare both panels). We also observed colocalization of retrograde signals with the sparse diffuse anterograde signals in temporal and cingulate cortex (cyan arrowheads). To quantify this reciprocity, we integrated the total amount of anterograde and retrograde tracer signals in each cortical area and plotted their correlations (Fig. 6G). The correlation was high (r=0.94) in frontal cortex (red dots), where strong columnar patches were abundant, and lower but still robust (r=0.76) when non-frontal areas (open circles) were included, where diffuse signals dominated. Similar results were found in the other 13 dual-injection cases, supporting the generality of these observations (Fig. 6H, Fig. S6).

### Corticostriatal projections also consist of patchy and diffuse projections

PFC has massive unidirectional projections to the striatum, constituting the first step of the cortico–basal ganglia–thalamo–cortical loop that plays an integrative role in goal-directed behaviors^35,36^. Anterograde tracer injections in the macaque suggest that the corticostriatal projections include both “focal” (patchy) and “diffuse” projections^37^. In marmosets, we also observed dense patches of anterograde label surrounded by sparser and more widespread anterograde label; here, we characterize these patterns in detail using an analysis strategy similar to that applied above to corticocortical projections. The striatum includes the caudate nucleus (Cd), putamen (Pu), nucleus accumbens (Ac), and tail of the caudate nucleus (Cdt), which are implicated in different functional circuits^36^. Typical termination patterns for three exemplar pairs of injections in these structures are shown using linear scaling (Fig. 7A) to emphasize the strong focal/patchy projections, and in log-scale view (Fig. 7B) to emphasize the diffuse spread of weaker projections. In the linear view, we observed multiple patches along the rostrocaudal axis mainly within the caudate nucleus, consistent with previous observations in macaques^38^. The patches were discrete for dlPFCd (#29, green) and dlPFCv (#42, red) injections (Fig. 7A, left panel) and were more distributed for other injections, particularly for the A25 injection (#81, yellow) that primarily targeted the nucleus accumbens (Fig. 7A; see also Supplementary Video 3). In the log-scale view, the tracer spread widely and there was extensive overlap across the different injections (Fig. 7B, lower panel). The patchiness of corticostriatal projections was evaluated by measuring the signal spread at a threshold of half the maximum value (Fig. 7C and 7D). Strong signals were concentrated in the caudate nucleus, except for the A25 injection (targeting nucleus accumbens) (Fig. 7D). The projections from dlPFCv were the most focal among the subregions, occupying less than one-seventh of that occupied by dmPFC or ACC injections. The location of these projections varied systematically within the caudate nucleus, with little overlap between adjacent subregions (Fig. 7E, Fig. S7C). As with the corticocortical projections, we could predict the centers of patches based on the injection coordinates using polynomial regression models (Fig. S7D-7G). These observations support the importance of topography in determining the specificity of focal (patchy) corticostriatal projections, consistent with previous studies in the macaque^37,39^. The axonal terminations included fine axonal fibers with bouton-like varicosities (Fig. S7A, B).

**Figure 7:**
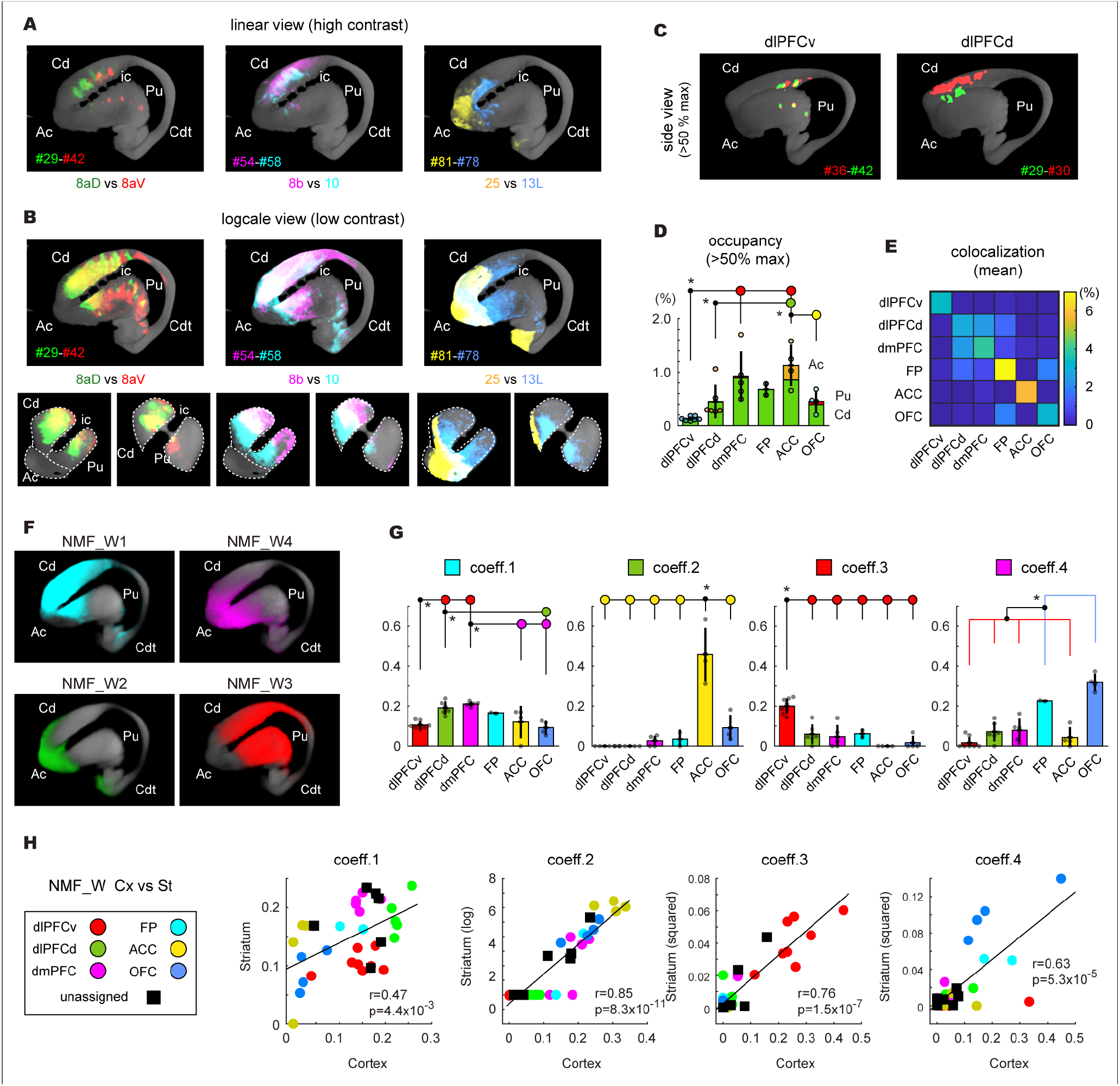
Characterization of focal (patchy) and diffuse corticostriatal projections. **A**, Three-dimensional (3D) representation of corticostriatal projections for three pairs of injections, shown with linear scaling, showing focal (patchy) axonal convergence. See Supplementary Movie S3 for bilateral rotation views. Ac, nucleus accumbens; Cd, caudate nucleus; Cdt, the tail of the caudate nucleus; Pu, putamen; ic, internal capsule. **B**, The log-scale view of the same data shows a diffuse spread of the tracer signals. Note that the contrasts of the intensity differences are high and low for the linear and log scale views, respectively. The lower panels show coronal section views at two AP positions. **C**, Interdigitation of focal projections visualized by binarization at 50% of the maximum values. **D**, Occupancies of the binarized signals in the total striatal volume. The averaged occupancies in the caudate, putamen, and nucleus accumbens are overlaid. **E**, Averaged co-localization ratio within and across subregions. See also Fig. S7C. **F**, NMF_W1 through W4 in 3D representation. **G**, Coefficients of different subregions for each component. **H**, Similarity of coeff. 1 through 4 between cortical and striatal projections shown by scatter plots. The values for striatal coefficients in the scatter plots show the original (for coeff. 1), log-transformed (for coeff. 2), or squared (for coeff. 3 and 4) values, which are correlated with the non-modified cortical coefficient values (see Spearman’s rank correlations and p-values in each panel).

To characterize the distribution of diffuse projections, we performed NMF analyses and generated four components (Fig. 7F). The NMF_W1 component, spanning most of Cd plus the anterior Pu, was strongly represented in all PFC subregions (Fig. 7G, coeff. 1), suggesting that the caudate nucleus is an important target for all the PFC subregions. Other striatal regions (e.g., Pu, Ac and Cdt) also received projections from some subregions of the PFC (compare Fig. 7F and 7G). Interestingly, the profiles of these coefficient value sets were similar to those of the corticocortical projections (Fig. 7H; Spearman’s rank correlation test). These observations strongly suggest that the global patterns of corticocortical and corticostriatal projections are governed by similar PFC topographic gradients, albeit in a nonlinearly skewed form. Visualization by the color indexing strategy confirmed that the PFC gradients were recapitulated globally in the striatum (Fig. S7H).

## Discussion

In this study, we report on the first quantitative analysis of anterograde tracer injections in marmoset PFC. Using a dataset with exceptionally high volumetric resolution and signal sensitivity, we defined and characterized two distinct projection types, patchy (columnar/focal) and diffuse, both of which contribute prominently to corticocortical and corticostriatal projections. Below, we discuss the topographic architecture of the primate brain and the functional implications of two types of projections, including the question of whether cortical columns in PFC constitute discrete, well-defined entities that largely or completely tile the prefrontal cortical sheet.

### Topographic gradients in the primate brain

We found that both patchy and diffuse projections recapitulate PFC topography in their local and global patterns (Fig. 8A). The patchy (columnar) projections in the cortex include parallel streams to extra-frontal association areas. The full PFC domain is represented in the cingulate and temporal fields, whereas the dorsolateral domain is emphasized in the parietal field. The diffuse projections from different PFC locations are widely distributed across the cortical hemisphere with large overlaps, reflecting the PFC gradients globally.

**Figure 8:**
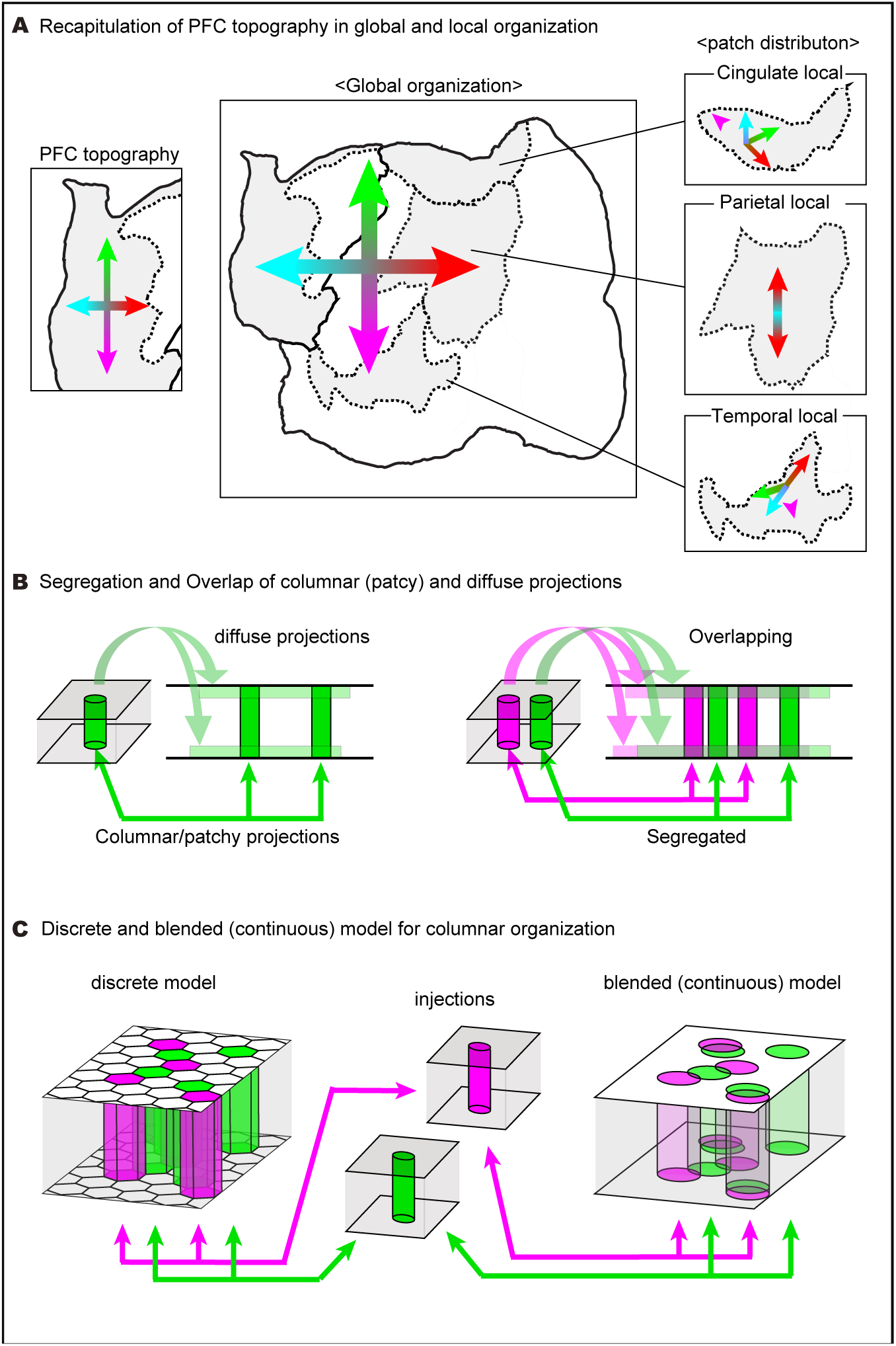
Summary of differential areal and lamina patterns of the columnar patches and diffuse projections. **A**, Schematic representation to show recapitulation of the PFC gradients in the global pattern of the diffuse projections and the local patterns of the columnar patch distributions. The cingulate and temporal fields each receive columnar projections from all the six PFC subregions, whereas the parietal field receives columnar projections only from the dorsolateral surface. Although skewed, the layout in each field resembles the layout in the PFC. **B**, Differential lamina preferences and overlaps of columnar and diffuse projections. Columnar projections more or less extend in the radial directions and converge into small areas, whereas the diffuse projections spread widely within the top and bottom layers. Because of such differential spread, the columnar projections are expected to segregate even from the nearby regions (right panel, green and magenta), whereas the diffuse projections show large overlaps in the target regions (mixed color). **C** Discrete vs blended (continuous) models for columnar organization. The discrete model postulates columnar modules with well-defined borders and no overlapping inputs and outputs with neighboring columns, whereas the blended (continuous model) postulates no such borders. If the former model is correct, any two remote injections would would always show complete overlap or complete segregation. Otherwise, partial overlap (blending) would be a common occurrence.

Topographic organization has been proposed as a key organizing principle for various PFC extrinsic projections^31,36,38,40–43^ and functional properties^44–46^. Recently, a mapping of the macaque lateral PFC by electrical microstimulation-fMRI reported mirroring of lateral PFC maps in five association cortex regions^23^. Our data suggest that such mirroring also occurs for at least three cortical regions in the marmoset, likely involves orbitomedial PFC as well, and also applies to corticostriatal connections. Precise correspondences between macaque and marmoset association areas have yet to be fully established, despite many cytoarchitectonic and connectional similarities^4,22^. The presence of gradients that are similar but not identical in the temporal, parietal, and cingulate fields provides important insights into both conserved and divergent connectional architecture across primate species. Regarding the global gradients, topographic similarities exist between the macroscale network organization in humans and anatomical connectivity in marmosets^30,47^. These observations suggest that the dual global/local topographic organization for the marmoset PFC projections may include features that are widely conserved across the primate lineage.

### Functional implications of PFC connectivity patterns

The concept of “parallel distributed networks” has been used to explain subdivision-specific connectivity patterns in the macaque PFC^42^ as well as the parallel interdigitated networks within carefully mapped individual human subjects^48^. This concept appears to apply to the patchy projections in our study, which connect distinct populations of source and target neurons reciprocally in the cortex (Fig. 8B, right panel, green and magenta columns). An intriguing question is whether each column in PFC is a discrete entity that has well-defined borders with no overlapping inputs and outputs with neighboring columns (Fig. 8C, left). Alternatively, a column identified by tracer injection might partially overlap and be “blended” with a column identified by a tracer injected in a different location (Fig. 8C, right). Our observation that anterograde and retrograde columnar patches were in most cases precisely coextensive (Fig. 6A-C) is consistent with the discrete columnar model but does not prove it. If further experiments indeed confirm the discrete columnar model for PFC, a host of fascinating questions arise: How many columns are in each PFC area? How many other columns does a given PFC column project to and receive inputs from? Are there gaps between neighboring columns, or is the PFC completely tiled by a mosaic of columns? How do neighboring columns differ in connectivity and function, gene expression patterns and/or cell type composition? Recent advances in anatomical tracer methods^49,50^, sub-millimeter fMRI^51^, and spatial transcriptomics^53^ will likely yield important insights regarding these issues.

A discrete columnar system in PFC would differ fundamentally from columnar systems in visual cortex that have been intensively studied, particularly in the macaque (see Introduction). An iso-orientation domain in V1 is in essence a thin ribbon that winds through the cortical sheet, but the ribbon lacks a well-defined thickness because orientation preferences change continuously rather than in discrete steps. An ocular dominance stripe in V1 has a finite thickness (width) related to eye-specific geniculocortical terminations, but along the length of a stripe, features such as orientation and spatial frequency appear to be mapped continuously^9,52^. Area V2 has a tripartite arrangement of thick stripes, thin stripes, and interstripes and putative columnar systems for representing multiple dimensions, including orientation, binocular disparity and hue (reviewed in^8,54^). In higher visual areas, modular organization has been reported in area V4 related to color and to shape and in inferotemporal cortex related to faces, bodies, color, disparity, and objects^10,55– 57^. However, we are not aware of compelling evidence in any extrastriate visual area for discrete columns at a sub-millimeter scale of the type hypothesized above for PFC and perhaps other higher association areas.

In contrast to the patchy PFC columnar system, diffuse projections spread widely and overlap extensively (Fig. 8B, mixed color area in the top and bottom layers). An important question is the degree to which anterograde signals in deep layers (the DL and DL2 patterns in Fig. 5C) represent fibers of passage rather than just axonal terminations. Our confocal microscopy observations revealed a low density of branches and boutons along with the sparse axon fibers in deep layers (Fig. S4E), suggesting these are not exclusively fibers of passage. In the dual tracer experiments, the deep-layer DL and DL2 domains are generally associated with sparse retrogradely labeled neurons, consistent with these regions being reciprocally connected (Fig. S6C). Thus, while we acknowledge that fibers of passage may be common, we consider it likely that most DL domains, and probably many DL2 domains as well involve sparse but functionally relevant reciprocal pathways. Since diffuse projections are typically weak, significant modulatory effects might arise mainly when a large population of neurons exhibits coordinated activity (Fig. S8, upper panel). This suggests that anatomical connectivity defines the broad framework of brain circuits and that their precise neural activity patterns depend on the dynamic state of the entire system. In this regard, resting-state fMRI in marmosets suggests the existence of such coactive networks involving frontal areas, including a candidate for the default mode network^15,18^. We propose that primate PFC is a system that has both modular and integrative properties, and its discoordination may contribute to mental disorders (Fig. S8). This raises the intriguing question of the degree to which diffuse vs patchy/columnar projections arise from and/or terminate on separate vs overlapping neuronal populations. Increasingly powerful molecular genetic, single-cell and/or ultrastructural approaches are likely to inform these questions.

Neural recordings from the monkey PFC suggest that computations in the PFC emerge from the concerted dynamics of large populations of neurons^58^, in which multidimensional activities may be superimposed^59,60^. Our findings suggest that the columnar/patchy and diffuse connectivity might serve as substrates for the segregation and integration of population activities, respectively. We find it notable that the laminar profile of projections commonly differ for the columnar and diffuse projections even in the same area (Fig. 8B). Laminar profiles have been considered to reflect the directionality of information flow and hence the hierarchical organization of cortical areas^7,32,34^. Our data suggest inconsistencies with a traditional hierarchical scheme for PFC organization, but the issues of laminar profiles and hierarchical organization will benefit from further analyses using more extensive datasets, which are currently ongoing. Another fundamental issue is the need for a more accurate cortical parcellation of marmoset PFC, incorporating multimodal data including gene expression data. Finally, we note that our marmoset PFC connectivity database offers many exciting opportunities to perform fine-grained analyses that were not previously possible and should contribute to our understanding of PFC structure and function in primates.

## Acknowledgments

We thank N. Hasegawa and RIKEN ARD/RRD for marmoset care. We thank Drs. Kathleen Rockland, Nenad Sestan and Takuya Hayashi for critical reading of the manuscript. We thank the RIKEN CBS-Olympus Collaboration Center for the technical assistance with confocal image acquisition. This work was supported by the program for Scientific Research on Innovative Areas (grant number, 22123009) from MEXT, Japan, by the program for Brain Mapping by Integrated Neuro technologies for Disease Studies (Brain/MINDS: JP15dm0207001to T.Y. and JP22dm0207088 to K.N.) from AMED, Japan and by NIH grant RO1 MH-60974 to D.C.V.E.. This work was made possible in part by software funded by the NIH: FluoRender: Visualization-Based and Interactive Analysis for Multichannel Microscopy Data, 1R01EB023947-01 and the National Institute of General Medical Sciences of the National Institutes of Health under grant numbers P41 GM103545 and R24 GM136986.

## Author contributions

Conceptualization, A.W., D.V.E. and T.Y.; Methodology, A.W., M.T., and J.H.; Software, A.W., H.S., F.R., Al.W., R.G; Investigation, A.W., J.H., and J.W.; Resources, A.W., M.T., H.M., H.S., F.R., Al.W, R.G.; Data Curation: A.W., H.S., Al.W.; Writing – Original Draft, A.W.; Writing –Review & Editing, A.W., H.A., K.N., N.I., H.S., D.V.E., H.O., S.I., T.Y.; Funding Acquisition, K.N., and T.Y.; Supervision, T.Y., S.I., and H.O.

## STAR Methods

### Marmoset experiments

All experimental procedures were carried out following the National Institute of Health Guide for the Care and Use of Laboratory Animals (NIH Publications No. 80-23) revised in 1996 and the Japanese Physiological Society’s “Guiding Principles for the Care and Use of Animals in the Field of Physiological Science,” and were approved by the Experimental Animal Committee of RIKEN (W2020-2-009(2)). The age and sex of the marmosets used are listed in Supplementary Table 1. We did not distinguish these factors in this study.

In our standard procedure, we performed MRI under anesthesia at least one week before surgery to plan injections. The presumed positions of cortical areas were determined by registration of the Brain/MINDS reference^22^ and the stereotaxic positions were aligned by using the interaural plane and the anterior end of the cortex. Surgery for tracer injections was performed as previously described with some modifications^61^. Pressure injection was performed using a glass micropipette with an outer diameter of 25–30 μm connected to a nanoliter 2000 injector with a Micro4 controller (World Precision Instruments). For exposed cortical areas, we injected 0.1 μl each of tracers at two depths (0.8 and 1.2 mm from the surface), aiming to deliver the AAV to all cortical layers. For deep injections (e.g., OFC), we injected 0.2 μl of tracer at one depth. With these volumes, we did not experience overflow, unlike the experience in our previous study^61^. To avoid fluorescence cross-talk, all subjects received a fluorescent tracer at a single site. However, in some of our samples, we injected nonfluorescent tracers, such as BDA, into several other locations, but these results are not reported here. After surgery, the marmosets were returned to the cages and sacrificed four weeks later.

### AAV tracers

We used a TET system to amplify fluorescence signals^62–65^. This system labeled all the known projections from the PFC and did not show strong cell-type bias. The detection of two types of projections is also not due to our TET labeling system, as we observed a similar convergence and spreading of axon fibers using the conventional biotinylated dextran amine (BDA) method (data not shown). Our standard tracer mix included AAV1-Thy1S-tTA (1 × 10e9 vg/μL), AAV1-TRE-clover (1 × 10e9 vg/μL; a GFP derivative), and AAV1-TRE3-Vamp2-mTFP1 (0.25 × 10e9 vg/μL; cyan fluorescent protein targeted to the pre-synapse). In later experiments, we also included AAV2retro EF1-Cre (1.5 × 10e9 vg/μL) in the tracer mix for co-injection. We deposited plasmids for these AAV vectors to Addgene (https://www.addgene.org/).

### PFC injections

In this study, we present the data from 44 high-quality datasets involving injections into various frontal areas of the left hemisphere (Fig. 1H, Fig. S1G, and Supplementary Table 1). To plan injections, we referred to previous studies^4,5,43,66,67^. We divided the frontal areas of marmoset into nine subregions, including the frontopolar cortex (FP), dorsolateral PFC_ventral (dlPFCv), dorsolateral PFC_dorsal (dlPFCd), dorsomedial PFC (dmPFC), ventrolateral PFC (vlPFC), anterior cingulate cortex (ACC), orbitofrontal cortex (OFC), as well as premotor areas (PM) and dorsal ACC (dACC) (Fig. S1G). The ACC subregion in this study comprises areas A32, A14, A25, and A24a and corresponds to the subgenual anterior cingulate cortex (sgACC) and perigenual ACC (pgACC) in other studies^68^ and was differentiated from the dACC comprising A24b and A24c. We injected most densely into dlPFC, which were further subdivided into “dlPFCv” and “dlPFCd,” corresponding to the areas 8aV/A45 and 8aD/46D, respectively, by our cortical annotation; dlPFCv most likely overlaps extensively with the frontal eye field (FEF)^30,69^. The relationship of “dlPFCd” to macaque areas 46, 9/46, and 8aD is currently unclear. However, our results suggest the inclusion of A46-like areas in dlPFCd, judging from the auditory projection (Fig. S3D). Throughout the manuscript, we used 29 datasets with clean injections in six PFC subregions (FP, dmPFC, dlPFCd, dlPFCv, ACC, and OFC) for subregion comparison. These injections are shown color-coded in Fig. 1H and Fig. S1G. Some of the injections localized at the border of these subregions and were used only for parcellation-free analyses (gray dots in Fig. 1H). Injections into the vlPFC, PM, and dACC subregions were also used only for the selected analyses.

### Serial two-photon tomography imaging (STPT)

After transcardial perfusion with 4% paraformaldehyde in 0.1 M phosphate buffer (pH 7.4), the marmoset brains were retrieved, post-fixed at 4 °C for 2−3 days, and transferred to 50 mM phosphate buffer (pH 7.4). All marmoset brains were subjected to *ex vivo* MRI before further processing. Agarose embedding was performed as previously described^28^ in 5 % agarose using a custom-made mold. Before embedding, we treated the brain with 1 mg/ml collagenase (Wako 031-17601) at 37 °C for one hour and manually removed the meninges as thoroughly as possible. This step ensured direct crosslinking of agarose to the brain for stable sectioning at a 50-μm interval. STPT was performed as described^26–28^ using TissueCyte1000 or TissueCyte1100 (TissueVision). We immersed the agarose block in 50 mM PB supplemented with 0.02 % sodium azide for imaging and sectioning. We used a 920-940 nm laser for excitation, and the two-photon excited fluorescence was recorded in three-color channels. Three-color imaging was critical in our study to (1) distinguish lipofuscin granules from tracers, which are characteristic of aged brain^70^ (Fig. S1F), and (2) to delineate AAV-infected cell bodies around the injection site, which was only possible in the unsaturated blue channel (ch3 Fig. S1D). In our setup, the resolutions for x and y were 1.385 μm/pixel and 1.339 μm/pixel, respectively. Both hemispheres were processed in the rostral-to-caudal direction (coronal). We imaged four optical planes (12 μm intervals) per 50 μm slice for the rostral part and one optical plane per slice for the caudal part. We used only the first optical plane in this study. One run consisted of 20−30 sessions with different XY coverage and took approximately 10 days to process the whole brain.

### Post-STPT histology

Sliced tissue sections were manually collected for further histological analysis. To confirm the cytoarchitecture, we collected one in ten sections for Nissl staining. The floating sections were mounted onto a glass slide after the agarose was removed. Mounted sections were rehydrated with PBS, and dark-field backlit images were taken using an all-in-one microscope (Keyence BZ-X710) before Nissl staining. The backlit image showed shading patterns that were very similar to conventional histological myelin stain (manuscript in preparation). We used the same section for Nissl staining, and the combined information of the backlit and Nissl-images provided useful information for anatomical delineation (Fig. S2). AAV2retro-Cre was detected by anti-cre recombinase (Milipore, clone 2D8), followed by Cy3-conjugated secondary antibody. The nuclear staining was imaged by an all-in-one microscope (Keyence BZ-X710) and processed for signal recognition.

For confocal microscopic observations, we stained the retrieved sections with the anti-GFP antibody (abcam, ab13790) and anti-Homer1 antibody (FRONTIER INSTITUTE, ner1-Rb-Af1000) in conjunction with Alexa488 conjugated anti-chick antibody and Cy3-conjugated anti-rabbit secondary antibody. After mounting on a glass slide, the stained sections were imaged using an Olympus Fluoview FV3000 confocal microscopy using a 40× silicone immersion lens (UPLSAPO40XS).

### Image processing pipeline

The details of the image processing pipeline are described elsewhere^24,71^. Briefly, the raw images in the three channels were stitched after background correction. The Ch1 (red) image provided the tissue background image with some tracer bleed-through, and the Ch2 (green) image provided the tracer fluorescence and tissue background. In our setup, we observed very weak signals for Ch3 (blue), which helped identify AAV-transduced cell bodies that were not identifiable with other channels due to signal saturation. Using this information, the pipeline automatically determined the exact spread of AAV transduction. The spread was calculated to be 2.6 ± 1.3 mm^3^ (mean ± SD) in the STPT template space (see below). The pipeline also accurately separated the axon signals from the background based on the Ch1 and Ch2 images. On visual inspection, we encountered virtually no false positives (e.g., lipofuscin granules misidentified as axons) for this process. Axonal detection was quite sensitive, although it was usually partial, meaning that only a part of visually recognizable axon fibers were labeled as the positive signals. The stitched coronal section slices (more than 600) were placed in the 3D space based on the recorded stage coordinates and non-linearly registered in 3D to the standard reference space or STPT template (see below). In this data transformation, a unit isocube (‘STPT-voxel’) corresponds to a 50 μm × 50 μm square of one slice image (50 μm interval), whose signal intensity was determined by the sum of positive pixels (out of ∼1400), each of which was weighted for fluorescent intensity by its 16-bit encoding. In this way, we secured a wide dynamic range of quantification and a high signal-to-noise ratio. All quantitation was based on the fluorescence-weighted axon signal values.

Before normalization (see below), the total sum of thus-defined signal intensities was highly variable between samples due to various experimental factors, including the variable spread of the tracer near the injection site. However, there was little correlation with age (r = -0.062; p = 0.69) or sex (p = 0.38, by a t-test), which we did not analyze further in the current study.

To evaluate the registration accuracy, we compared the registered Ch1 images of individual samples with the STPT template. More precisely, we determined the boundary positions separating the cortex, putamen, globus pallidus, and internal capsule for each sample along a line ROI on horizontally sliced images (Fig. S1A). These boundaries were determined automatically by selecting the peaks of optical density changes near each target boundary (red arrows (i)-(v)). As shown in the whisker plot, the deviance was kept within a few STPT voxel units (corresponding to 50-μm isocubes). Although this evaluation does not guarantee accurate registration for all the voxel points of the entire brain, it demonstrates highly accurate registration for regions near high-contrast landmarks, such as the white matter.

### Standard reference space (STPT template)

To facilitate 3D-3D registration between optically-based and MRI-based volumes, we built a standard reference image for a marmoset brain based on iteratively averaged Ch1 images^24^. We call this 50 μm-isocubic 3D image as the “STPT template.” To allow multimodal data integration, the initial averaging step was performed using BMA2019 Ex Vivo (Space 2), which is a 100 μm isocubic reference image based on a population average of 25 *ex vivo* MRI (T2) contrasts^72^. Nissl-based cortical annotation originally performed on a single brain^22,73^ was transferred to the STPT template and used without further adjustment.

### Handling of 3D data for visualization and substructure analyses

After completing the pipeline, the tracer intensity data and AAV-transduced area data were converted to the STPT template-registered form. These 3D data were converted to a NIFTI format (https://nifti.nimh.nih.gov/), and structures of interest could be easily excised for each sample using the labelmap. The labelmap for the definition of substructures was manually annotated based on the image contrast of the STPT template (Fig. S1B) using 3D Slicer^74^ [https://www.slicer.org/]. After excising the structures of interest, the tracer intensities were standardized to the maximum value within the substructure, converted to 8-bit or 16-bit, and saved as a tiff stack file. FluoRender^75^ was used to visualize 3D data for the presentation of images and movies. Striatal data sometimes included strong contaminating signals of passing axon fibers within the internal capsule and white matter. In such cases, we manually masked these regions using a 3D Slicer (https://www.slicer.org/) and performed the normalization again.

### Conversion of cortex data into a flatmap stack and its normalization

The outer (pial) and inner (white matter) segmentation surfaces of the cerebral cortex were defined based on the image intensity of the STPT template and manually corrected when necessary (Fig. S1B). A 167,082-vertex surface mesh from the STPT template with vertices approximately midway through the cortical sheet provided an initial geometric substrate for generating pial, white, and midthickness surfaces that were in topological correspondence with one another. Radial trajectories determined by a dense orientation field created by a heat propagation model were used to generate the pial surface via vertex migration to the outer cortical boundary, the white surface via vertex migration to the inner cortical boundary, and the midthickness surface via migration to the midpoint of the radial trajectory linking the pial and white vertices. Fifty equally spaced vertices were identified along each radial trajectory connecting the inner and outer surface vertices (Fig. S1C, right hemisphere). The Brain/Minds cortical midthickness flatmap^72^ based on the same 167,082-vertex mesh as the above 3D surface was used to define the x-y coordinates of a flatmap representation; the z dimension was represented by the spacing between vertices along each radial trajectory in the 3D volume. ANTs nonlinear registration^76^ was then used to generate a deformation field that aligned the 50-layer array of vertices in the 3D cortical model to the corresponding set of vertices in the flatmap stack. The deformed cortical volume was then resampled using trilinear interpolation to generate a flatmap stack volume based on a 500 × 500 × 50 array of ‘flatmap-stack-voxels’ (fm-vox; Fig. S1C, left hemisphere)^24,71,72^.

To evaluate the relationship between tangential distances in the 3D model vs the flamap stack, we placed seed points at the midthickness layer in the flamap stack with 50 fm-vox spacings. These seed points were mapped to the STPT template space using the deformation field. Spheres of 15-STPT-vox radius were placed at each seed point in the STPT template space and then mapped back to the flatmap stack using the deformation field (Fig. S1E). The mean cross-section area was 247 fm-vox^2^. Given that the cross-sectional area for the 15-STPT-vox sphere can be approximated by a circle with 15-STPT-vox radius (706.5 STPT-vox^2^), and one STPT-vox corresponds to 50-μm in the STPT template space, one fm-vox in the flamap stack based on these values is on average 84 μm on a side.

To standardize the intensities of the cortico-cortical projection patterns of the flatmap stack, we prioritized the constancy of signals in the upper layers (levels 26−50) because the lower layers contained more diffuse and widespread signals, some of which may be passing fibers. We also knew that the signal levels of the tracers at the injection site are unreliable due to fluorescence saturation. Therefore, we first averaged layer levels 26−50 of the flatmap stack to make a 2D flatmap in which columnar convergence could be detected as peaks of fluorescence intensity. Herein, one pixel corresponds to tangential components of one fm-voxel. We masked the injection area based on the MIP of all layers of the flatmap stack for the injection site segmentation, because fluorescence intensity is usually saturated and not reliable in such area. We then selected pixels with the top 0.1% of the intensity values from the rest out of those that constitute the 2D flatmap and set the minimum value of the selected pixels at 10,000. This standardization method provided very similar patterns across different intensity ranges within the same frontal group irrespective of the original tracer intensity. The same coefficient was used to standardize the flatmap stack. For the six-color overlay of different PFC groups in Fig. 1I, the MIP images for layer level 26−50 from the same PFC subregions were MIP-merged to generate a 2D flatmap. The flatmaps of different groups were then overlaid with different colors in a “winner-take-all” style, in which the color of the strongest signals was displayed to avoid color mixing. For the six-color overlay in Fig. S1H, the color was simply merged when overlapped.

To convert the tracer intensity data into a log-scale representation, the standardized values were log-transformed after adding 0.001 and thresholded by 0. For visualization, the log-transformed data were scaled to an 8-bit representation. We routinely used the Videen color palette (obtained from Connectome Workbench color palette; https://github.com/Washington-University/workbench) for pseudocolor intensity scaling.

### Detection and characterization of columnar patches in corticocortical projections

Columnar patches were detected as local maxima in the 2D flatmap generated by averaging the upper layers (layer levels 26-50) of the 3D flatmap stack, with the injection sites masked and the signal values standardized as described above. Local maxima in 2D were identified using the “find peaks” function of MATLAB 2019a, inspired by the Fast 2D peak finder^77^ (https://www.mathworks.com/matlabcentral/fileexchange/37388-fast-2d-peak-finder), MATLAB Central File Exchange. Briefly, the 2D image was Gaussian smoothed, and peaks larger than the defined minimum height values were searched in the x- and y-directions, pixel by pixel. Only 2D spots that peaked in both the x-and y-direction searches were identified as local maxima. The minimum height value for columnar patch detection was set at 100, i.e., two orders of magnitude lower than the standardization value of 10,000. Columnar patches detected at the fringe of the injection site due to masking were removed manually by visual inspection. We detected 1867 columnar patches from 44 datasets.

To examine the extent of convergence of tracer signals to the centers of columnar patches, we set three 3D ROIs for measurement (Fig. 2C); namely, central cylinder with a 4-fm-vox radius, a 19×19×50 fm-vox rectangular cuboid (hereafter referred to as 19-vox cuboid) and a 50×50×50 fm-vox cubic (hereafter referred to 50-vox cubic). The idea is to measure the ratio of the number of tracer-positive voxels contained within the cylinder to that in the 19-vox cuboid and the 50-vox cube. First, we standardized the intensity of the tracer signals by the maximum value within the upper half of the central cylinder. Second, the tracer signals were binarized at 0.5, half the maximum value. Third, the largest clump of the binarized tracer signals within the upper half of the central cylinder was chosen to represent the columnar patch of choice (Fig. 2B and 2C, green “clump”). The ratio of the number of voxels within the central cylinder to that of all voxels that constitute this binary clump within the 19-vox cuboid and 50-vox cube was calculated as the index of central convergence. Columnar patches were often clustered, and stronger tracer signals were at nearby locations connected to the central clump. Therefore, we classified the columnar patches into the “solitary” and “connected” groups based on the degree of central convergence. When we set 50% of central convergence for the 50-vox cubic ROI as the threshold, 1008 patches out of 1867 patches fulfilled this criterion. The central convergence for this group of patches was equal to that of the 19-vox cuboid ROI, which means that the segmented central clumps of the tracer signals were contained within the 19-vox cuboid and disconnected from the surrounding signals. Hence, we called this group “solitary”. The patches that did not fulfill this criterion were called “connected” because the central clumps for over 97 % of the remaining 859 patches stretched out of the 19-vox cuboid. Because of the relatively high degree of central convergence, the morphology of the central clumps for the solitary groups could be reasonably well approximated by ellipsoids whose major axis lengths, centroid positions, and angles were estimated by regionprop3 function (CY Y; 2022; https://github.com/joe-of-all-trades/regionprops3).

### Comparison of mouse and marmoset data for columnar projections

All the mouse data used in this study were obtained from Mouse Brain Connectivity Atlas^26,27^ (https://connectivity.brain-map.org/). First, we selected the datasets with injections into the frontal areas (ACAd, ACAv, FRP, ILA, MOs, ORBI, ORBm, ORBvl, PL) of the wild-type mice (Supplementary Table 3). The whole-brain tracer data (projection energy) with 10 μm resolution for these datasets were then acquired via the application programming interface (API) and converted to flatmaps as previously described^78^. The flatmaps for both hemispheres were resized to 816×408 pixels. The image intensity was standardized similarly to that used for the marmoset flatmap except that the injection center was not excised before measurement. The columnar patches were detected using the same code used for the marmoset flatmap. The mouse flatmaps represented the average of all layers, whereas the marmoset analysis used flatmaps representing only the upper half of the layers. To compare with the mouse data, we performed a new patch detection using the flatmaps of all layers for comparison (Fig. 3E and 3G).

Our detection algorithm simply finds the peaks above a threshold in the X and Y directions after smoothing. It can therefore detect broad peaks that do not necessarily extend radially like those of the marmoset cortex. It is also sensitive to standardization methods. The mouse data differ from our marmoset data in several ways. (1) The mouse study used regular AAV-GFP, which is less efficient at labeling axons than our TET-enhanced AAV. (2) Mouse brains were sliced at 100 μm intervals as opposed to 50 μm intervals in our data, which is less reliable in detecting submillimeter scale structures. (3) The frontal region of the mouse is curved, and flatmapping causes large distortions. (4) We could not subtract the tracer signals of the cell bodies before standardization for the mouse data. All these differences could lead to differences in the detection of the columnar patches in the two species. However, the original slice images suggest that the projection patterns of axons are very different between mice and marmosets (e.g., Fig. 3B and 3F). Therefore, the quantitative estimation in Fig. 3G is considered legitimate overall.

### Prediction of columnar patch distribution patterns by polynomial regression for corticocortical projections

To examine the topographic projections to the cingulate and temporal fields, we set the ROIs for these areas, as shown in Fig. 2G, and determined the center of mass of the columnar patches present within these ROIs for each sample for the six PFC subregions. Among the 29 injections, 24 had at least one columnar patch in the cingulate field, and 21 had columnar patches in the temporal field. To find a regression model to predict the projection from the injection, we used a polynomial regression of degree 2, which exhibited a better corrected AIC score than degree 3. As predictor variables, we used the X and Y coordinates of the injection center in the flatmap. As response variables, we used the X or Y coordinates of the averaged positions of columnar patches in the cingulate and temporal ROIs. As shown in Fig. S3A and 3b, we achieved reasonable fitting (R^2^ =0.68∼0.92) for the injections in the PFC. The permutation test, in which the models constructed by the original and shuffled data were compared for accurate reconstitutions, confirmed the effectiveness of the regression model. To extrapolate the obtained topography, we calculated the projections from every point in the PFC map to the cingulate and temporal fields, as shown in Fig. 2H. This calculation resulted in projections outside the flatmap, suggesting the presence of distortions at the fringes. Fig. 2H displays only the projections within the flatmap region.

### Non-negative matrix factorization (NMF) analyses of corticocortical and corticostriatal projection patterns

To find common patterns of projections for corticocortical (or corticostriatal) projection, we performed NMF analyses using log-transformed data. NMF has been used as an effective method of dimensionality reduction for neuroimaging^79^. Unlike principal component analysis, all coefficients and basis images are nonnegative, allowing a more straightforward interpretation of the data^80^. For corticocortical projections, 500 × 500 fm-vox MIPs of all layers with injection site masks were downsized to 100 × 100 size, and the data for 44 injections were converted to a 44 × 10,000 matrix to be used for “nnmf” function of MATLAB. For corticostriatal projections, a 180 × 300 × 220 STPT-vox space containing the left striatum was downsized to a 45 × 75 × 55 space, and a 44 × 185,625 matrix was used for NMF analyses. NMF analysis attempts to decompose the original matrix A (sample × pattern) into basis images (or components) W and coefficients H (A ≅ WH). Due to the nonnegative constraints, A is not exactly equal to WH. Furthermore, the nnmf function of MATLAB uses an iterative algorithm starting with random initial values for W and H, and the results obtained vary each time slightly. To find the best result, we repeated the analyses 100 times and examined the residuals between A and WH. The residuals were relatively constant but sometimes became large. Inspection of the basis images showed that very similar images were generated when the residuals were small. Therefore, we selected W and H with the smallest residuals as the basis images. Because we had only 44 datasets for analysis, we tried to minimize the number of basis images while achieving adequate reconstitution efficiency. With four basis images, the averaged correlation coefficients between the reconstituted and the original images were approximately 0.927 (Fig. S4B). Reconstitution by five basis images did not result in substantial improvement (0.934) and we chose to use four basis images for further analysis.

Conversion of NMF_W1, 2, 3, and 4 into a single colormap was performed by first combining the W1/W4 and W3/W2 pairs by subtraction. By this subtraction, W1 (or W3)-dominant regions take positive values, whereas the W4 (or W2)-dominant regions take negative values because of the nonnegative feature. Furthermore, because of the antagonistic relationship of these pairs, there exist relatively small overlaps that cancel each other’s value. As a result, the positional information of the four components (or basis images) was mostly retained in the positive and negative domains of the subtracted values (compare Fig. 4B with Fig. 4D). Using the 2D colormap shown in Fig. 4G^81^, these two value sets can be jointly represented by color-coding. Note that this color-coded map has no information about the intensity and just the relative ratio. In showing this map, the region with the low contribution of any of the four components was excluded. Conversion of coefficients 1, 2, 3, and 4 into a colored dot map was performed similarly.

### Analysis of retrograde labeling

We used AAV2retro-EF1-cre as the non-fluorescent retrograde tracer. This construct accumulates CRE protein in the nucleus, which results in relatively even labeling of variously sized neurons. It was also advantageous for the automatic identification of labeled cells. After immunohistological detection and imaging, the retrieved image data were processed for automated segmentation of labeled nuclei and registration to the corresponding STPT image (FR and HS, manuscript in preparation). Although we collected only one in ten sections to detect the retrograde tracer, we were able to map the result of retrograde tracing to the STPT template by registering the slice images to the STPT slice data. To evaluate the colocalization of anterograde and retrograde tracers at the columnar scale, we measured the image intensity by “Plot Profile” function of FIJI using the vertical line ROI shown in Fig. 6B^82^ [https://imagej.net/software/fiji/]. We also measured the amount of each tracer in the 117 cortical areas of the flatmap stack format for the examination of correlations. The correlation coefficients in Fig. 6G and Fig. S6 were calculated based on the log10 of the original signal values. We did not adjust the signal values for the size of each area and we excluded areas with fewer than threshold (10^0.5^=∼3.2) retrograde signals from calculations of correlation coefficients.

Visual inspection of the antibody-stained images showed that some injection cases showed a low signal-to-noise ratio, as judged by an abundance of false-positively detected artifactual signals across cortical areas. We distinguished such samples by calculating the ratio of the numbers of areas with low values against those above the threshold values, excluding areas with no signals. The samples that failed this quality check exhibited lower correlation coefficients (Fig. S6B), suggesting that high signal-to-noise ratio of the retrograde tracing data, which is determined by the sensitivity and specificity of immunolabeling affects correlation with the anterograde data. In two cases, we also observed substantial leak of the retrograde AAV into the white matter, which appeared to have transduced the cells via passing fibers. We excluded these two cases from further analyses.

Although the anterograde and retrograde tracer distributions were well correlated overall, there were mismatches of various kinds. These mismatches are partly due to purely technical confounds and partly to differences in methodology. Technical confounds included gaps in data arising from the wider interval between sections analyzed for retrograde data or misregistration, especially in distorted regions on the fringes. In addition to these technical reasons, methodological differences could lead to mismatches. One significant difference is that the retrogradely labeled cell is counted as one, irrespective of the amount of tracer incorporated from the axon terminals. For this reason, we suspect that the diffuse connectivity is less efficiently labeled in the retrograde labeling and that its identification depends greatly on experimental conditions. On the other hand, our anterograde tracing cannot distinguish the passing fibers from the synaptic connections and may overestimate genuine “connectivity”, especially for sparse signals. Thus, the evaluation of reciprocity using the dual tracer system that we adopted in this study requires careful consideration of each case.

### Lamina-profiling of the columnar and diffuse cortical projections

Inspection of signal distributions for individual columnar patches indicated various laminar preferences for these patches. Because the directions of the axonal extension were not always perpendicular to the flatmap but were largely contained within the central cylinder of 4-fm-vox radius, we measured the maximum intensity of tracer signals within the central cylinder for each of the 50 layers. We defined it as the laminar profile of the columnar patch of interest. We performed hierarchical clustering to classify the laminar profiles of the 1867 columnar patches. Pearson’s correlation coefficient was used to measure distance, and Ward’s method was used for the clustering. Although we selected columnar patches based on the values of layer level 26−50, the profiles of all 50 layers were used to calculate the correlation coefficient. Fig. 5A shows the original laminar profiles of the columnar patches (before averaging) aligned according to the tree structure of the hierarchical cluster analysis. We divided them into eight clusters (separated by white border lines) and averaged each of the laminar profiles shown in Fig. 5A. We further defined three lamina types, “DL,” “ML”, and “UL”. The defined lamina types were color-coded blue, green, and red, respectively, for representation in flatmap format in Fig. S5B.

To examine the laminar profiles of the overall areal projections, including both the columnar and diffuse projections, we divided the cortical hemisphere into 117 areas and examined the averaged laminar profile in each area. When we calculated the sum of tracer signals for the 44×117 injection-to-target area pairs, 97% of the pairs had non-zero signals. To assess the significance of these signals, we aligned the log10 values of the sum of the signals in order and found that the decline in log10 values gradually accelerated around the 4,000^th^ pair and dropped sharply after the ∼4,500^th^ pair. Based on this observation, we decided to use the top 4,000 injection-to-target area pairs for laminar profiling as the pairs with significant connections. The hierarchical clustering was performed in a similar manner to that used for the focal projections, except that we made a new group called “DL2”, which showed very restricted signal distribution near the gray/white matter interface. There is a possibility that some of these signals stem from the intrusion of white matter fibers of passage. At present, we cannot distinguish between synaptic contacts and fibers of passage, although we did observe bouton-like varicosities to be associated with apparently passing-fiber-like signals by confocal microscopy (Fig. S4E).

### Occupancy and colocalization of focal (patchy) tracer signals in the striatal region

We noticed that the occupancy of tracer signals in the striatal regions varied widely among the PFC subregions when focal/patchy projections were visualized (Fig. 7A, linear view). To quantitatively estimate this point, we set the threshold at 50% of the maximum intensity (Fig. 7C) and counted the ratio of positive voxels in the caudate nucleus, putamen, and nucleus accumbens, separately over the entire striatal voxels for each sample. The values in the three striatal compartments were summed for analysis by ANOVA to compare the six PFC subregions. For the colocalization analysis, the ratio of the overlapping voxel numbers to the number of combined voxels was calculated for every pair of 29 injections (Fig. S7C). Due to patchy distributions, overlaps within the same PFC subregions were not necessarily high and varied greatly among pairs.

### Detection of patches of corticostriatal projections in 3D

For the detection of patches of corticostriatal projections in 3D, the 3D data were first converted to a set of XY and YZ MIP images for the detection of local maxima in each 2D image in a similar way used for the detection of corticocortical columnar patches, and the 3D positions that fit both MIP images were selected as the local maxima in 3D. When two patches were detected within a six-STPT vox distance in 3D, we combined them for simplicity. The detected patches were projected onto the XY, YZ, and XZ MIP images for visual inspection for accurate detection. In the detection of caudate patches, we found that strong signals in the Muratov bundle or internal capsule were sometimes included and selected as patches. In such cases, we either masked those regions for re-standardization or simply deselected them. Tracer intensities were standardized to the maximum values within the striatal region for each sample, and the minimum height value for patch detection was set at half the maximum value. An example of such patch detection is shown in Fig. S7D. The distribution of these patches could generally be well approximated by an elongated ellipsoid (Fig. S7D, red ellipses), consistent with the previous observation that macaque corticostriatal projections are longitudinally aligned^38^.

### Prediction of corticostriatal patch distribution patterns by polynomial regression model

To find regression models that can predict the projection coordinates from the injection coordinates, we tested the polynomial regression model with degree 2 and searched for the optimal fit. As predictor variables, we used the x, y, and z coordinates of injections in the STPT template space. As response variables, we used the x, y, or z coordinates for the average positions of the detected STPT-vox intervals that roughly correspond to AP = +13.5, +14.5, +15.5, +16.5, and +17.5 in the Paxinos atlas and visualized the corresponding lines in the caudate.

#### Data and code availability

All the section image data of STPT, standardized 3D data for the whole brain, and the flatmap stack data for the cortical signals of the marmosets have been deposited at Brain/MINDS data portal (https://dataportal.brainminds.jp/marmoset-tracer-injection) and are publicly available as of the date of publication. The high-resolution images used in the paper are labeled with the marmoset number and the section number (e.g., #82-139) and are available through the zoomable section viewer. In addition to the visualization and search tools available at these sites, users can download standardized 3D data for tracer segmentation, fluorescence-weighted segmentation, AAV-transduced cell segmentation, and original and standardized flatmap stack data in nifti format on request. The mouse analyses used the existing, publicly available data of the Mouse Connectivity atlas. The accession numbers for the used datasets are listed Supplementary Table 3. Other Microscopy data reported in this paper will be shared by the lead contact upon request.

Any additional information required to reanalyze the data reported in this paper is available from the corresponding authors upon request

## QUANTIFICATION AND STATISTICAL ANALYSIS

### Statistical tests

We generally used 8, 6, 5, 2, 4, and 4 samples for dlPFCv, dlPFCd, dmPFC, FP. ACC, and OFC, respectively, for subregion comparisons. The injections that were positioned near the borders of these subregions were excluded in such cases. Values are reported as mean ± standard deviation (SD) throughout the manuscript. The correlation coefficients (r) in Figs 6G, 6H, Fig. S4B and Fig. S6 refer to the Pearson correlation. Those in Fig. 7H refer to Spearman’s rank correlation. R^2^ for the regression model refers to the coefficients of determination in Fig. S3A/3B, 3I, 4C, and 7E. The p-values were calculated using one-way ANOVA and Tukey’s post hoc test for significant factors in the ANOVA for Figs 2E, 4C, 7D, and 7G.

## Supplementary Figures

## Supplementary Tables

## Supplementary Videos

Supplementary Video 1: Columnar cortical projections from the marmoset PFC Three-dimensional presentation of the cortical projections from a PFC injection (A8aD) in the standard reference space. White matter signals are shown in red.

Supplementary Video 2: Contralateral columnar projection in 3D Three-dimensional reconstitution of the contralateral projections at high resolution.

Supplementary Video 3: Focal (patchy) and diffuse Corticostriatal projections Three-dimensional reconstitution of the corticostriatal projections. Focal (patchy) and diffuse projections are visualized in turn for representative samples.

## Supplementary Figs and legends

**Fig. S1:**
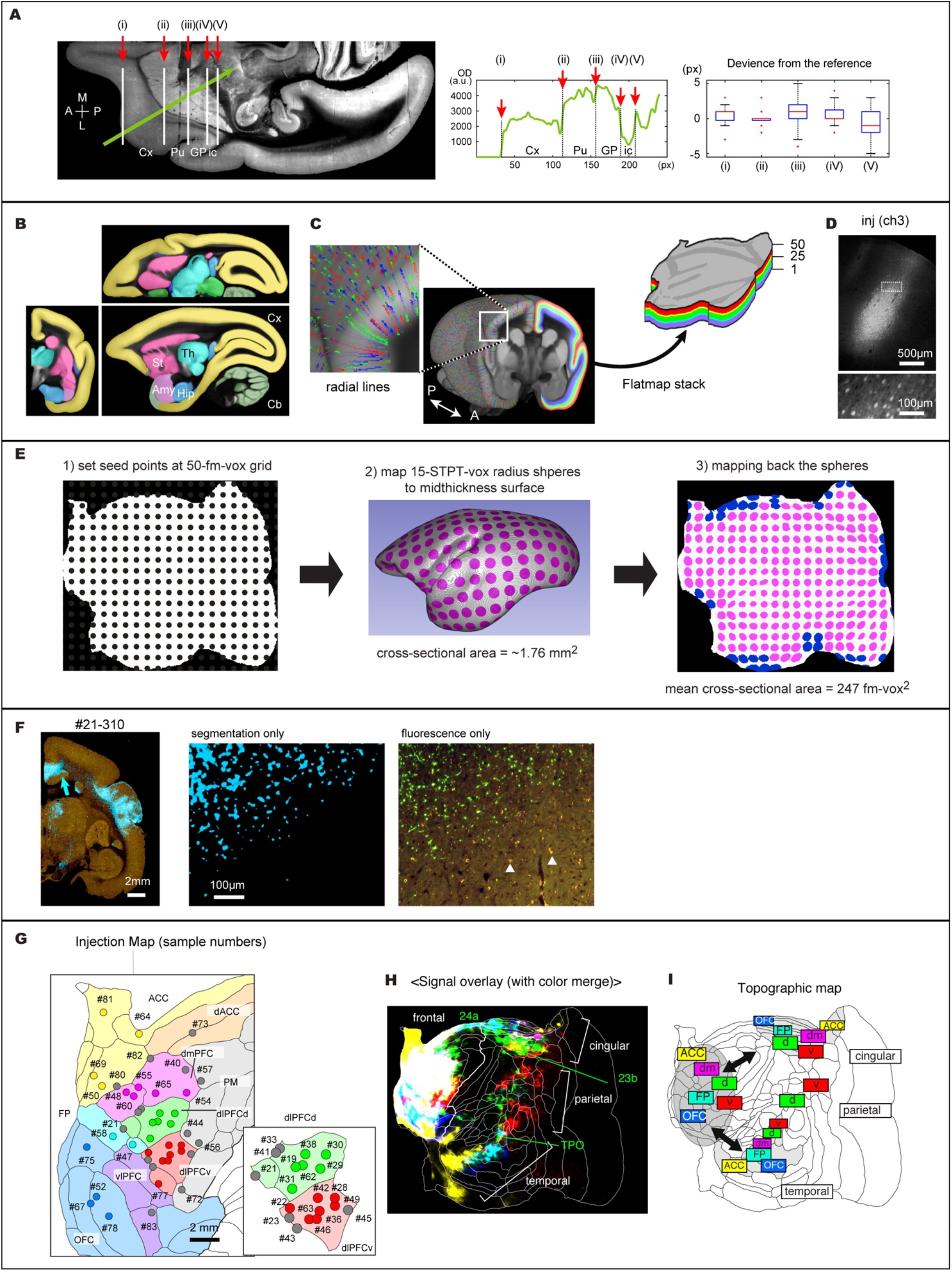
Mapping of PFC projections by serial two-photon tomography imaging (STPT). **A**, An example image showing how we estimated registration accuracy along one dimension (green line). The 3D-reconstructed images were registered to the STPT template and virtually sliced in the horizontal plane. Only the right hemisphere is shown (A; anterior, P; posterior, M; medial, L; lateral). The image intensity (optical density; OD) of individual samples was measured along the line ROI, which revealed the anatomical borders (i)-(v) separating the cortex (Cx), putamen (Pu), globus pallidus (GP), and internal capsule (ic). The deviation from each border determined for the STPT template is shown by box and whisker plots on the right. **B**, The section views for the STPT template overlaid with anatomical annotations. Cx, cortex; St, striatum; Th, thalamus; Amy, amygdala; Hip, hippocampus; Cb, cerebellum. **C**, Schematic sparse representation of the radial lines that determined the columnar structures of the cortical surface in different colors (right hemisphere) and the lamina structures determined by these vertices (left hemisphere). This information was used to convert the cortical surface into a stack of 50 layer-specific flatmaps. **D**, The bleed-through fluorescence in the blue channel used to locate the exact site of injection [inj(ch3)]. Note that the fluorescence of individual cells can be recognized unlike in saturated channels 1 (red) and 2 (green) (see Fig. 1A and 1C). **E**, A schematic explanation for how we measured the tangential intracortical distance in the flatmap stack. We first placed a grid of seed points on the midthickness layer in the flatmap stack at interval of 50 flatmap-stack-voxels (fm-vox). These seed points were mapped to the 3D STPT template space and spheres with 15-STPT-voxel radius were centered at each seed point. These spheres were mapped back to the flatmap stack using the deformation field. Finally, cross-sections of the deformed 15-voxel sphere in the mid-layer flatmap were used to calculate the approximate correspondence of 15-voxel distance in the flatmap stack. Any spheres that intersected the outer boundary of the flatmap or to an adjacent sphere were excluded from calculation (shown by blue color). **F**, Comparison of tracer segmentation (cyan) and original fluorescence. Aged marmoset brain sections typically contain many lipofuscin granules with dot-like autofluorescence^70^ (white arrowheads in the right panel), which was efficiently excluded in our signal detection algorithm. **G**, Estimated locations of injection sites in relation to six PFC subregions: dlPFCd (dorsolateral PFC-dorsal), green; dlPFCv (dorsolateral PFC-ventral), red; dmPFC (dorsomedial PFC), magenta; FP (frontopolar cortex), cyan; ACC (anterior cingular cortex), yellow; OFC (orbitofrontal cortex), blue. Others include vlPFC (ventrolateral PFC), dACC (dorsal anterior cingulate cortex), and PM (premotor areas), in which injection points are shown by gray dots. These colored injections were used as core samples for various analyses. Depending on the analyses, gray injections located outside the six PFC subregions (e.g., premotor areas) or on the borders between subregions were also used. See Supplementary Table 1. The uncertainty in localization in the tangential domain (parallel to the cortical sheet) is difficult to determine, but is likely to be within one mm, considering the projection patterns of each sample. The scale bar shows the approximate distance on the midpoint flatmap based on panel E. **H**, The overlay of projection patterns for six PFC subregions that allows color mixing. The convergence of projections from different subregions in A24a, A23b, and TPO, as well as most frontal areas, resulted in color mixing. Same as in Fig. 1I, except for color mixing. **I**, A schematic representation of the topographic projections deduced from the overlay of tracer signals for the six PFC subregions shown in Fig. 1I. d; dlPFCd, v;dlPFCv, dm; dmPFC.

**Fig. S2:**
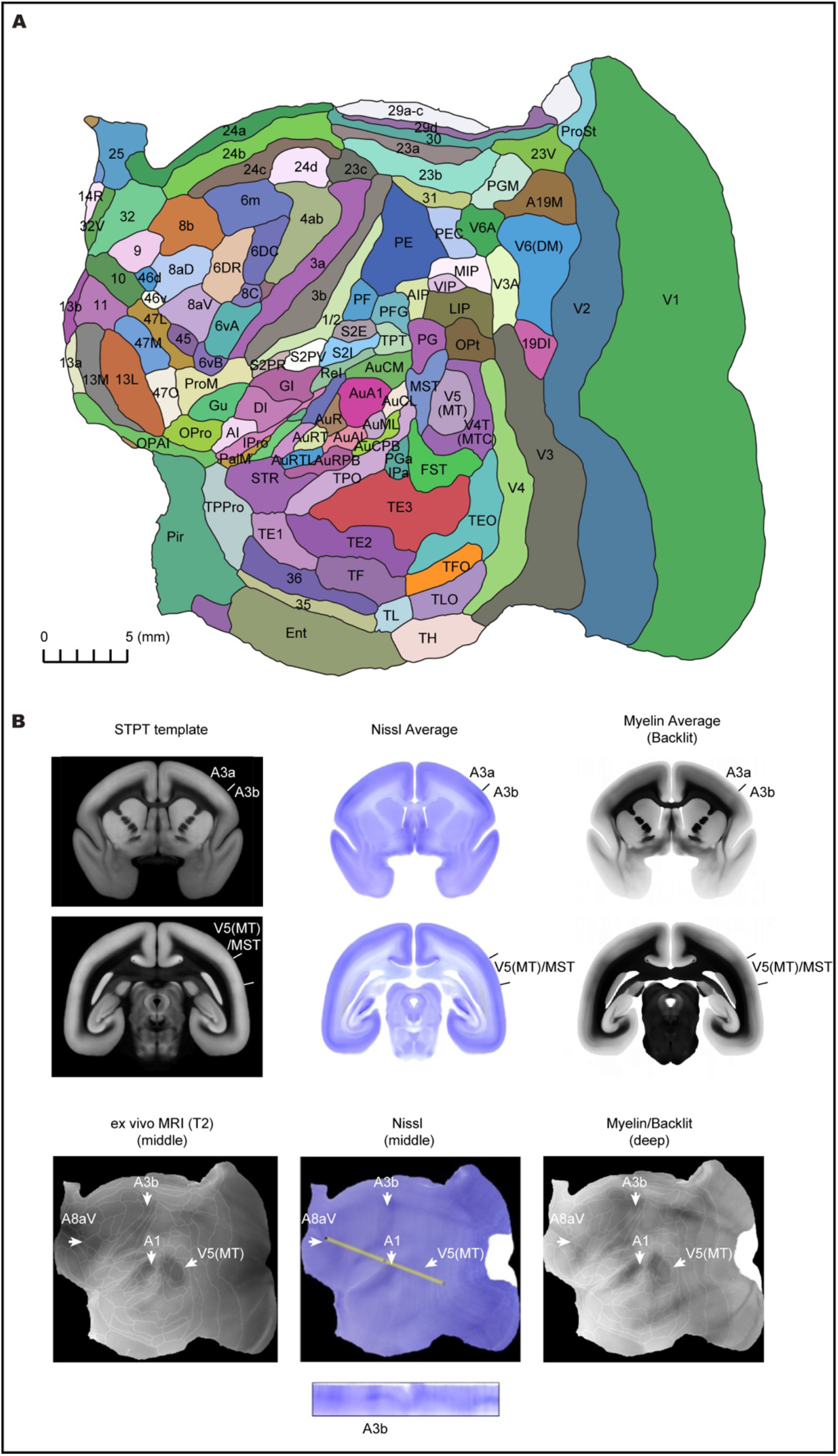
A map of the cortical areas and comparison with histological data. **A**, The cortical area annotation of the Brain/MINDS atlas^22,73^ is transferred to the standard reference space (STPT template) and the flatmap stack for registration. The scale bar shows the approximate distance on the midpoint flatmap based on Fig. S1E. **B**, Comparison of the cortical annotation of the STPT template with histological data. After STPT imaging, one in ten sections were retrieved for Nissl staining and backlit myelin imaging (see STAR Methods). Here, the section and flatmap views for the averaged Nissl and backlit (myelin) images are shown. Several conspicuous landmarks in these images corresponded well to cortical annotations.

**Fig. S3:**
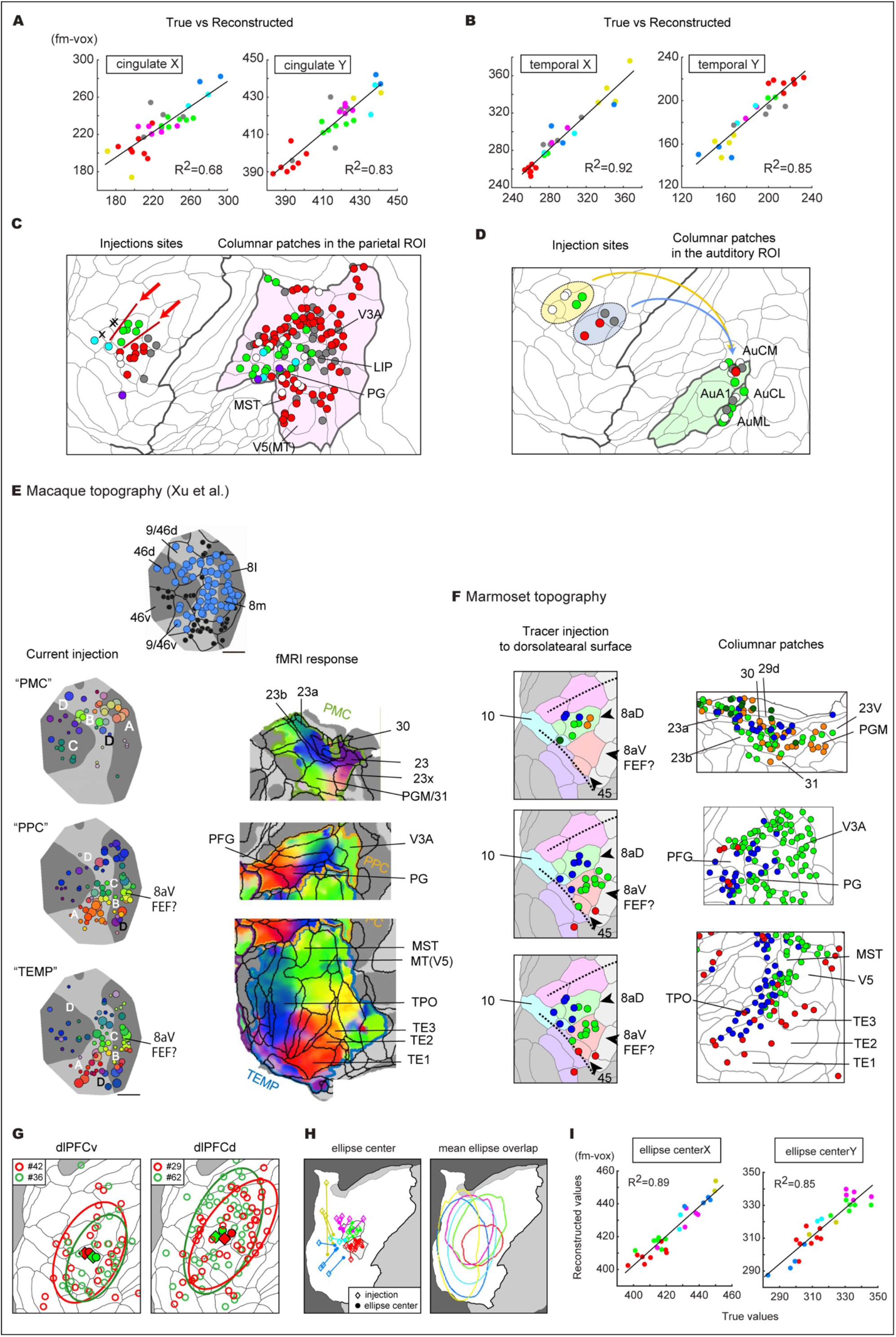
Topographic organization of the columnar patch distribution. **A, B**, Comparison of true and reconstructed values from polynomial regression models for topographic projections. The colored dots represent the injections into six PFC subregions, and the gray dots represent the border injections. **C**, Seed-based analyses to determine injections that exhibited columnar projections toward the parietal field. The colors of the injection sites and the columnar patches in the target ROI indicate which subregion they belong to (FP, cyan; dlPFCd, green; dlPFCv, red; vlPFC; purple; PM, gray; white, border). The red arrows indicate borders of abrupt changes in the projection profiles. The injections indicated by X did not generate columnar patches in the parietal field, demonstrating an abrupt border. **D**, Seed-based analyses to determine injections that exhibited columnar projections toward the early auditory field (core and belt regions). Projections from the anterior and posterior PFC regions intermingled in the auditory field, unlike the parietal field. **E**, Connectivity mapping of the macaque lateral PFC by stimulation-fMRI, reproduced from Xu et al (2022) with permission. Stimulation of the colored dots in the lateral PFC generates responses in the target regions with the same colors. Nomenclature for the cortical areas is as reported in Xu et al. **F**, Tracer injections in the marmoset PFC generate columnar projections with topographic relationships similar to the macaque counterparts. Injections into the colored dots generate columnar patches with the same colors in the target regions. Nomenclature for the cortical areas is as shown in Fig. S2. Note that the colors used for this panel are changed from other figures to highlight similarity with the macaque data. **G**, A pair of nearby injection sites in dlPFCv (left panel) and a different pair of nearby injection sites in dlPFCd (right panel). Note scattering of abundant columnar patches around the injection site, whose overall extent we approximated by ellipses enclosing 70 % of the patches. The injection sites (filled circles) and ellipse centers (diamonds) were slightly offset, but the topographic relationship between the two adjacent injections was preserved. The spread of patches was more restricted for dlPFCv than dlPFCd. **H**, (Left panel) The offsets between the injection sites (diamonds) and the ellipse centers (filled circles) for each sample of the six core PFC subregions are shown. These patterns reflect the wide dispersion of columnar patches from peripherally-located ACC and OFC injection sites (yellow and blue traces). (Right panel) The regions enclosed by colored lines represent regions occupied by the ellipses of at least half of the samples (>49 %) of each subregion. The overlaps of ellipses for each subregion were smallest for dlPFCv (red trace). **I**, Polynomial regression to fit the positions of the ellipse centers based on the injection coordinates.

**Fig. S4:**
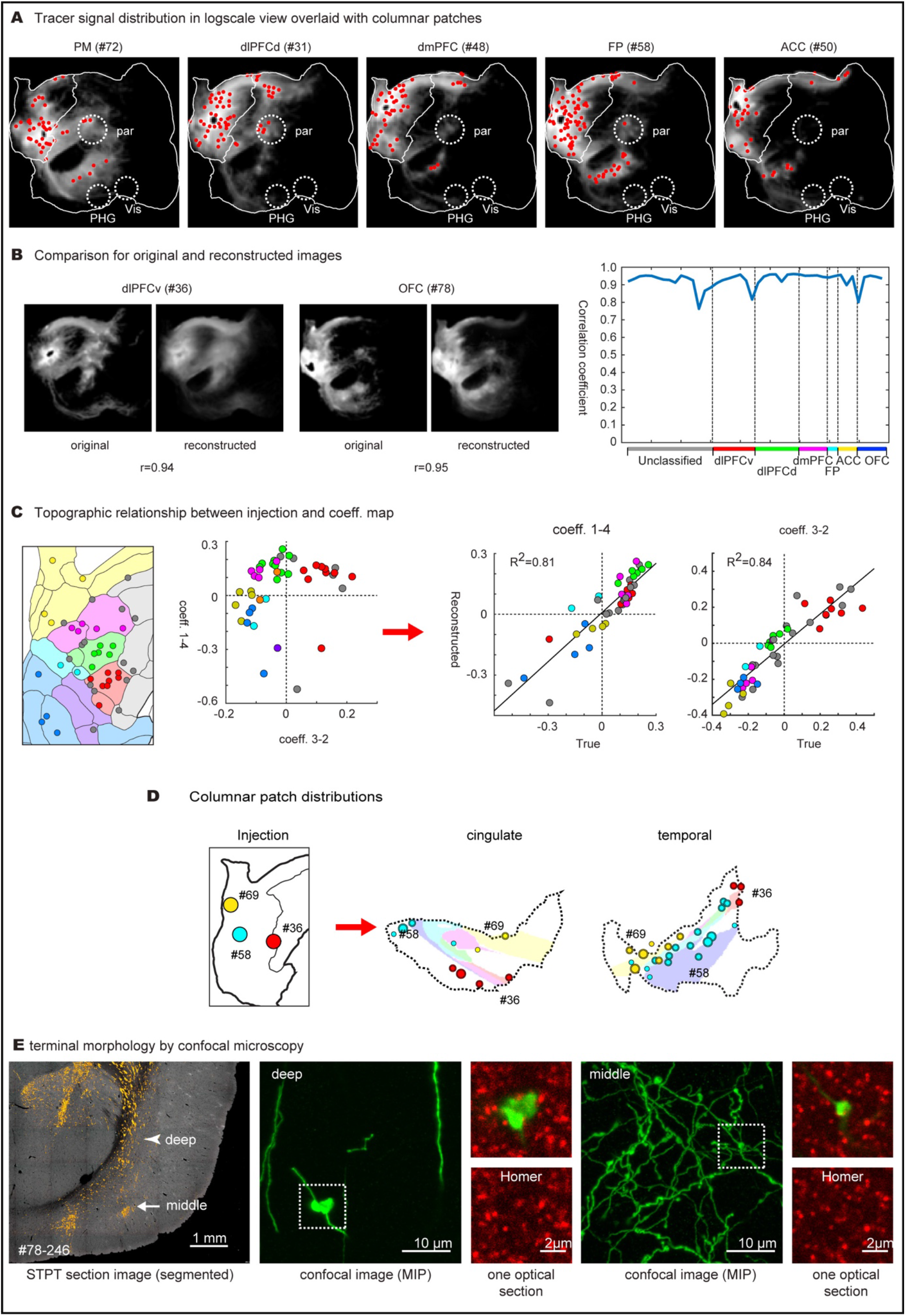
Characterization of diffuse corticocortical projections in log-scale view. **A**, Log-transformed tracer images from various frontal subregions, including premotor areas (PM), with detected columnar patches overlaid (red dots). **B**, Comparison of the original images and those reconstructed from the four NMF components. The accuracy of the reconstruction was measured using the correlation coefficients of the two images for 44 pairs. **C**, Comparison of topographic relationships of injection sites with plot positions in the coeff. 1-4 vs coeff. 3-2 value map. The polynomial regression models (degree 3) could predict these coefficients based on injection coordinates as shown. **D**, Global gradients affect the patch numbers and intensity for the columnar projections. Three example injections into dlPFCv (#36), FP (#58), and ACC (#69) generated columnar patches both in the cingulate and temporal association fields in a topographic manner, but the numbers and intensity greatly differed. The size of the circles represents the normalized intensity of each patch. **E**, Morphological examination of tracer signals by confocal microscopy. Tracer signals are examined in the deep (arrowhead) and middle (arrow) layers. The corresponding sections were retrieved for staining with anti-GFP and anti-Homer antibodies (a postsynaptic marker). The deep layer (arrowhead) and middle layer (arrow) regions imaged by confocal microscopy are shown.

**Fig. S5:**
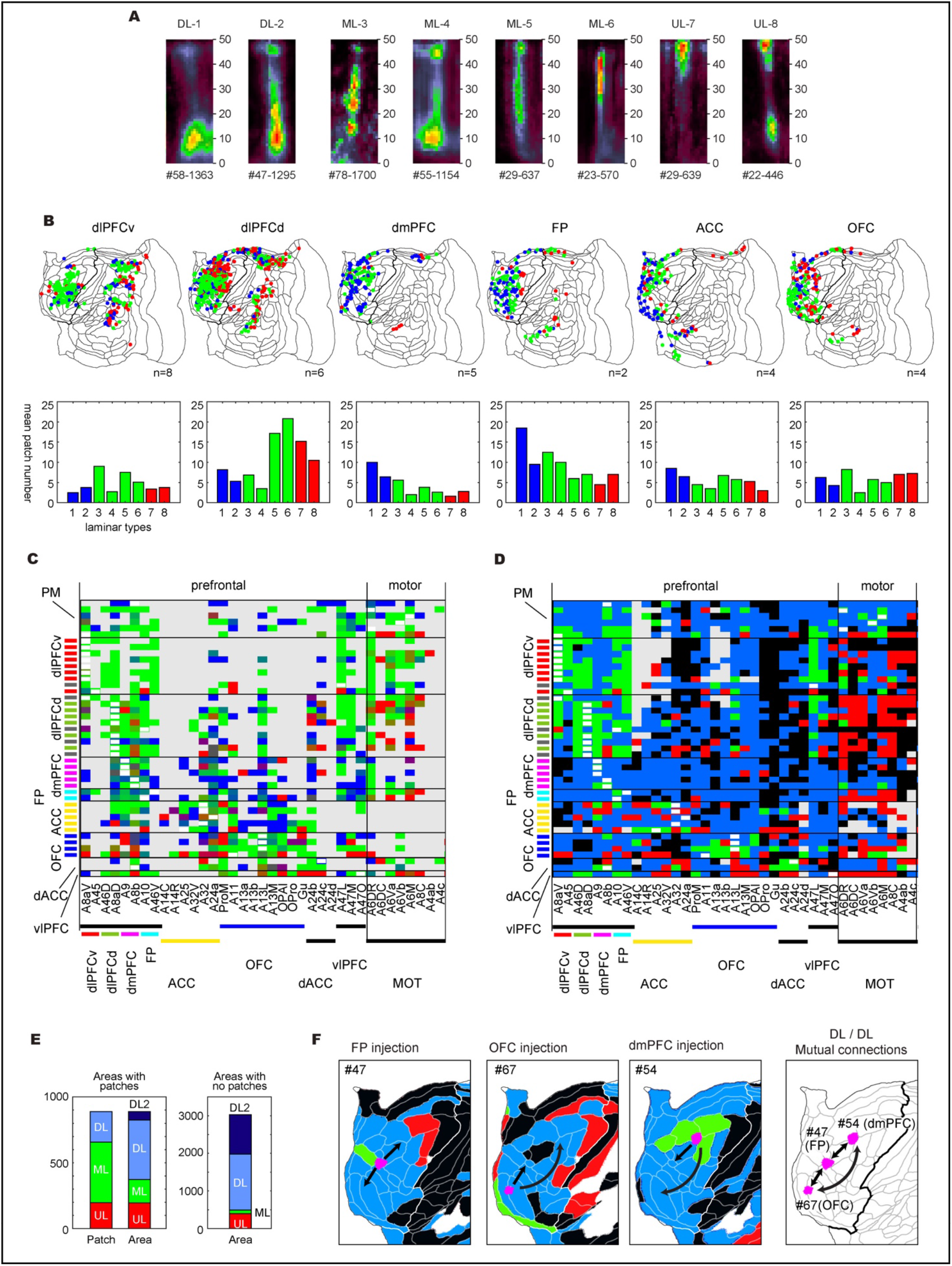
Laminar analyses of the columnar patches and the area-averaged diffuse projections. **A**, Representative laminar patterns for the eight clusters shown in Fig. 5A. **B**, Areal distribution of three lamina types (DL: blue, ML: green, UL: red) for each PFC subregion and the bar graphs for eight clusters. The colors of the bar graphs indicate the DL, ML, and UL types. The areal distribution shows the overlay of all samples, and the bar graphs show the mean values per injection. **C**, Ratiometric summary diagram showing the dominant lamina types for the columnar patches in 44×37 source-target combinations. The rows indicate each of the 44 injections and the columns indicate 37 cortical areas in the frontal cortex grouped into 9 blocks of similar localization (see Supplementary Table 2). The bars on the left side indicate injections into the six core PFC subregions. The gray bar indicates border injections. The areas of injections are shown by white rectangles. **D**, A summary diagram showing the area-averaged lamina types for the 44×37 source-target combinations. **E**, (Left panel) Comparison of dominant lamina types in the areas having patches with the area-averaged lamina types. (Right panel) Area-averaged lamina types in the areas with no columnar patches. **F**, An example of area-wise reciprocal connections showing DL types in both directions. A part of area-averaged lamina maps was shown for each of the three injections into FP (#47), OFC (#67), and dmPFC (#54).

**Fig. S6:**
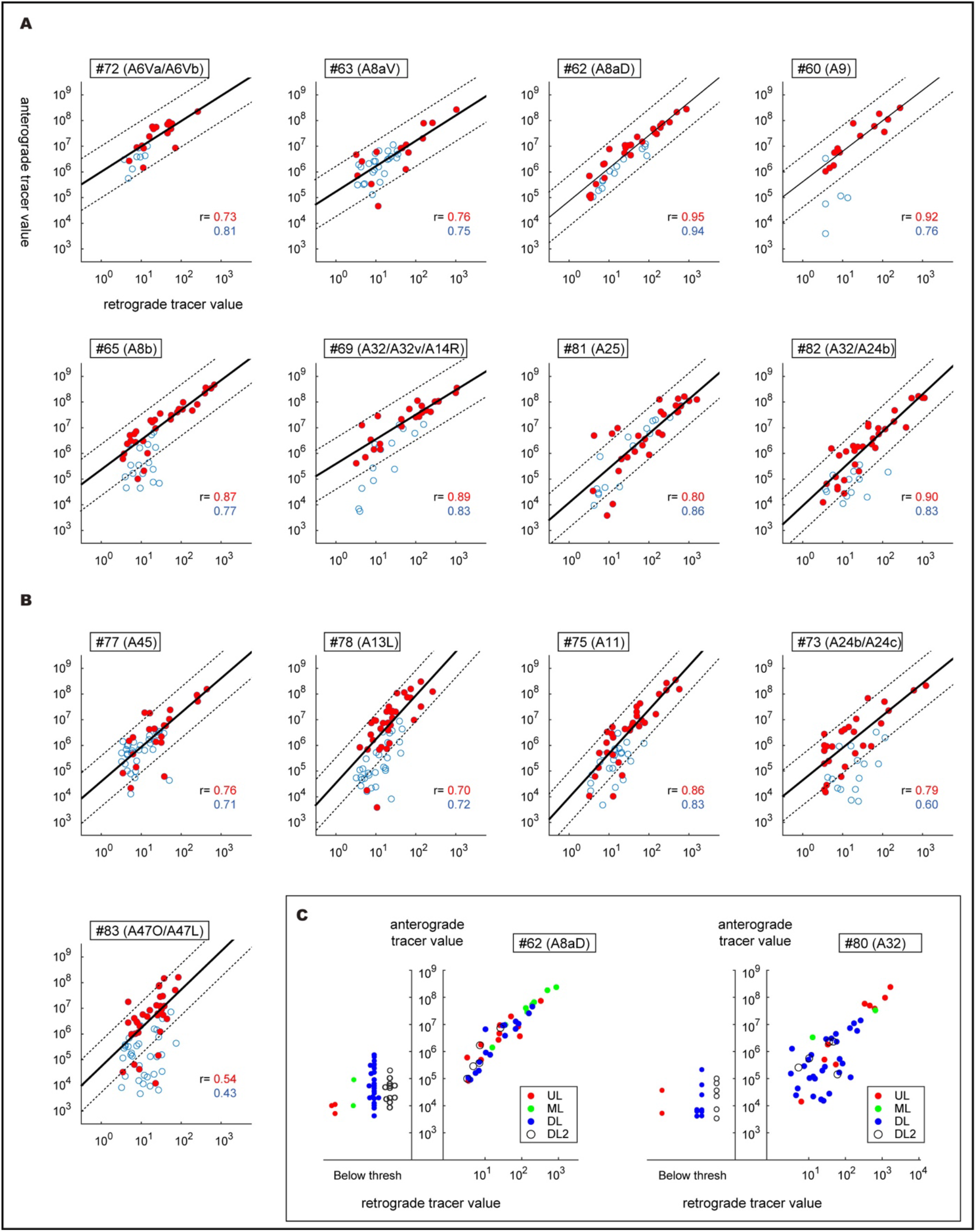
Area-based comparison of anterograde and retrograde signals measured in log scale. **A**, Samples with high-quality retrograde signals are shown (see method for the criteria). **B**, Samples with noisy retrograde signals. These samples had scattered signals throughout the cortical areas that appeared likely to be false positives based on visual inspection of the morphology of the signals. We still observed moderate signal correlations, especially for the strong signals. The retrograde signals for #83 largely consisted of noise artifacts by visual inspection. **C**, Two examples of anterograde-retrograde correlation in each area with laminar types assigned. Unlike in panels A and B and Fig. 6G, areas with below-threshold retrograde signals are indicated on the left side of the panel. As these examples show, DL (blue dot) and DL2 (open circle) types tended to be associated with low anterograde and retrograde signals.

**Fig. S7:**
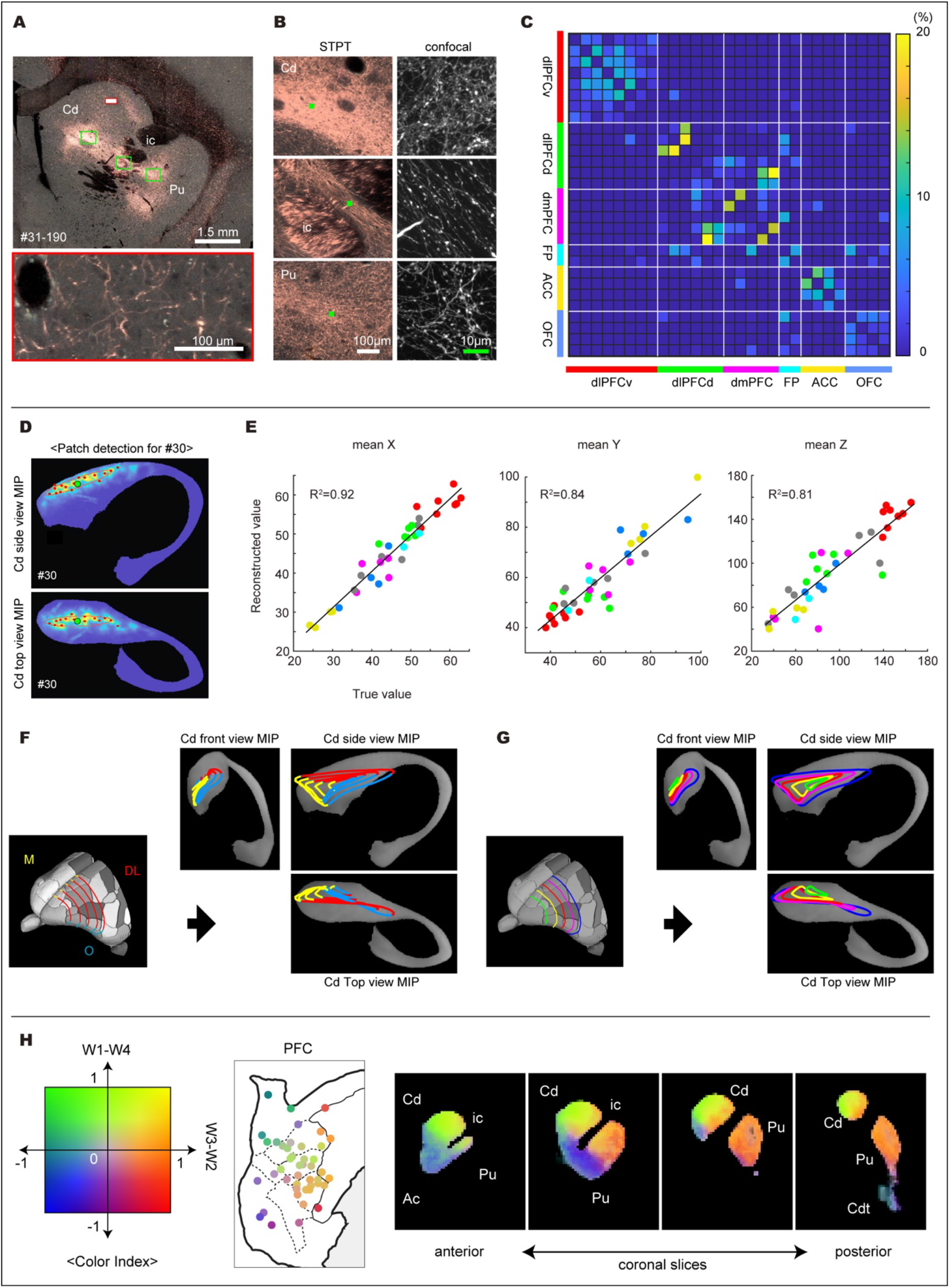
Focal (patchy) and diffuse corticostriatal projections. **A**, An example of the original section image of the corticostriatal projections. Here, the fluorescence of the tracer signals is colored pink. The white rectangle with red contours is magnified below as an example of a sparsely innervated region. The three green rectangles are magnified on panel B. **B**, A clump of dense tracer signals in Cd, a bridge region in the internal capsule, and a dense region in Pu are shown magnified. The green-filled rectangles in the STPT images indicate the approximate positions for confocal microscopy on the right panels. **C**, The colocalization of binarized tracer signals between two samples within and across six PFC subregions. See Fig. 7C for example images, and Fig. 7E for the averaged values. The average rate of colocalization was generally higher within the same PFC subregion than between different subregions (Fig. 7E), except that some dlPFCd samples showed similar distributions to dmPFC samples. Overall, however, the colocalization rate was low even in the same subregions. **D**, Detection of local maxima of tracer convergence foci; the average position of these foci is indicated by a black circle. The MIP side view and top view of the left caudate nucleus are shown. **E**, Scatter plots comparing the true and reconstructed values from polynomial regression models based on injection coordinates. Six core PFC subregions and border injections are used for regression fitting. **F**, Topographic projections from the frontal areas to the caudate nucleus predicted by the polynomial regression model. Five contours covering the dorsolateral (DL; red), orbital (O; blue), and medial (M; yellow) sides of the frontal cortex are projected onto the caudate nucleus. **G**, The five contours colored green, yellow, orange, red, and blue from the rostral to the caudal end are projected onto the caudate nucleus. **H**, NMF components and coefficients of the diffuse projections were color-coded in a similar manner to that used for Fig. 4H. The coronal sections roughly correspond to AP+12.5, AP+11.5, AP+9.0, and AP+7.0 in the Paxinos atlas (Majka et al., 2020) (https://www.marmosetbrain.org/reference). Ac, nucleus accumbens; Cd, caudate nucleus; Cdt, tail of caudate nucleus; Pu, putamen; ic, internal capsule.

**Fig. S8:**
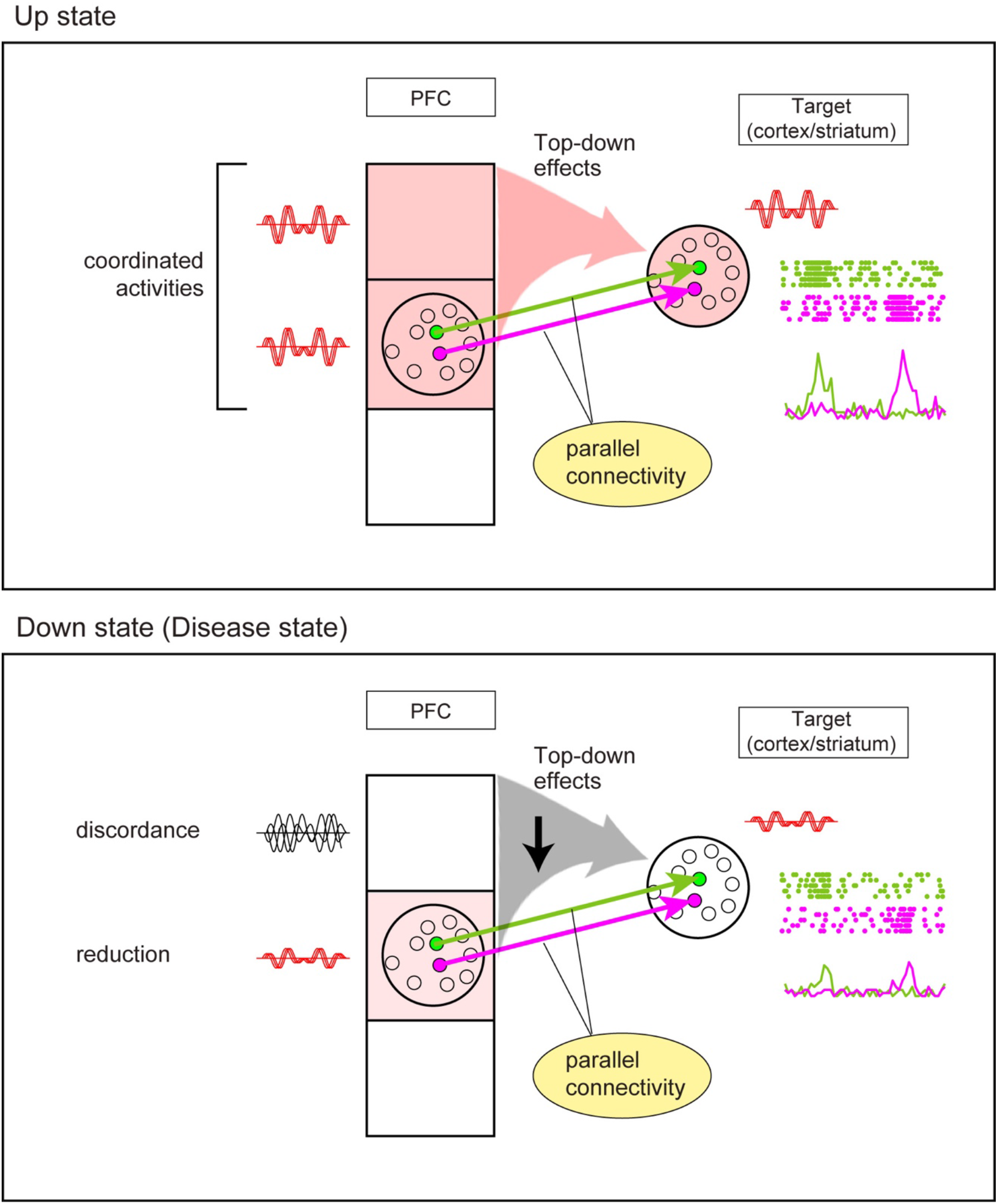
A model for state-dependent effects of the patchy and diffuse projections. [Up state] The patchy (columnar) projections involve a small population of neurons in a narrow space and may allow parallel information transfer (green and magenta arrows). On the other hand, the diffuse projections have overlapping distribution patterns (shown by a large arrow). If many neurons in the source exhibit coordinated activities, they could have significant modulatory effects on the target regions (cortex and the striatum). [Down state (Disease state)] Due to the low efficacy, the modulatory effects of the diffuse projections would be low or absent, if the source neurons lack coordination. It is hypothesized that psychotic symptoms of schizophrenia might be related to a reduced influence of prior beliefs on sensory data in Bayesian inference^83^. Such an “influence” might be encoded by the coordinated activities of PFC neurons conveyed via the diffuse projections (Upper panel, red arrow, “Top-down effects”). A disease state might arise from chronic reduction or discordance in such population-coded PFC activities (lower panel). In support of this idea, a change in functional connectivity networks has been suggested in Alzheimer’s disease and schizophrenia^84–86^.

**Supplementary Table1.** Marmoset injections analyzed in the present study.

| Data ID | Brain/MINDS ID | age (y) | sex | subregion | area | Core 6 PFC |
| --- | --- | --- | --- | --- | --- | --- |
| #40 | R01_0061 | 8.5 | M | PM/dmPFC | A8b/A6DM |  |
| #56 | R01_0083 | 6 | F | PM | A6DR |  |
| #57 | R04_0079 | 4.3 | F | PM | A6DM |  |
| #72 | R01_0104 | 5.3 | M | PM | A6Va/A6Vb |  |
| #44 | R01_0069 | 6.6 | F | PM | A6DR |  |
| #45 | R01_0070 | 6.6 | F | PM/dlPFCv | A8aV/A6Va |  |
| #49 | R01_0076 | 8.9 | M | dlPFCv | A8aV | o |
| #63 | R01_0091 | 2.4 | M | dlPFCv | A8aV | o |
| #36 | R01_0057 | 4.1 | F | dlPFCv | A8aV | o |
| #42 | R01_0063 | 8.3 | M | dlPFCv | A8aV | o |
| #46 | R01_0075 | 5.6 | F | dlPFCv | A8aV | o |
| #22 | R01_0029 | 5 | F | dlPFCv | A8aV | o |
| #28 | R01_0040 | 5.6 | M | dlPFCv | A8aV | o |
| #43 | R01_0064 | 8.3 | M | dlPFCv/vlPFC | A8aV/A47L |  |
| #77 | R01_0110 | 6.7 | F | dlPFCv | A45 | o |
| #23 | R01_0030 | 5.3 | F | vlPFC | A47L/A47M |  |
| #29 | R01_0043 | 3.3 | F | dlPFCd | A8aD | o |
| #31 | R01_0048 | 2.4 | F | dlPFCd | A8aD | o |
| #62 | R01_0088 | 3.7 | F | dlPFCd | A8aD | o |
| #19 | R01_0026 | 5 | F | dlPFCd | A8aD | o |
| #21 | R01_0028 | 5.1 | M | dlPFCd/FP | A10/A46D |  |
| #30 | R01_0046 | 3.4 | M | dlPFCd | A8aD | o |
| #33 | R01_0054 | 8.1 | F | dlPFCd/dmPFC | A8aD/A9/A8b |  |
| #38 | R01_0059 | 8.1 | M | dlPFCd | A8aD | o |
| #41 | R01_0062 | 8.5 | M | dlPFCd/dmPFC | A8aD/A9 |  |
| #48 | R01_0071 | 7.8 | M | dmPFC | A9 | o |
| #60 | R01_0094 | 10.8 | M | dmPFC | A9 | o |
| #55 | R01_0080 | 7.4 | F | dmPFC | A9/A8b | o |
| #65 | R04_0095 | 2.6 | F | dmPFC | A8b | o |
| #54 | R01_0081 | 7.6 | F | dmPFC | A8b | o |
| #47 | R01_0052 | 6.2 | F | FP | A46V/A10/A47L | o |
| #58 | R04_0080 | 3.2 | F | FP | A10 | o |
| #50 | R01_0072 | 10.4 | F | ACC | A32 | o |
| #80 | R01_0112 | 6 | F | ACC | A32/A9 |  |
| #64 | R01_0092 | 2.4 | M | ACC | A24a | o |
| #69 | R01_0095 | 2.3 | M | ACC | A32/A32v/A14R | o |
| #81 | R01_0114 | 9.8 | F | ACC | A25 | o |
| #52 | R01_0078 | 8.9 | F | OFC | A13L/A13M | o |
| #67 | R01_0090 | 2.9 | F | OFC | A13M/A13L | o |
| #78 | R01_0107 | 2.5 | M | OFC | A13L | o |
| #75 | R04_0058 | 5 | F | OFC | A11 | o |
| #73 | R01_0098 | 3.3 | M | dACC | A24b/A24c |  |
| #82 | R01_0115 | 8.4 | F | dACC | A32/A24b |  |
| #83 | R04_0023 | 5.2 | M | vlPFC | A47O/A47L |  |

**Supplementary Table2-1.**
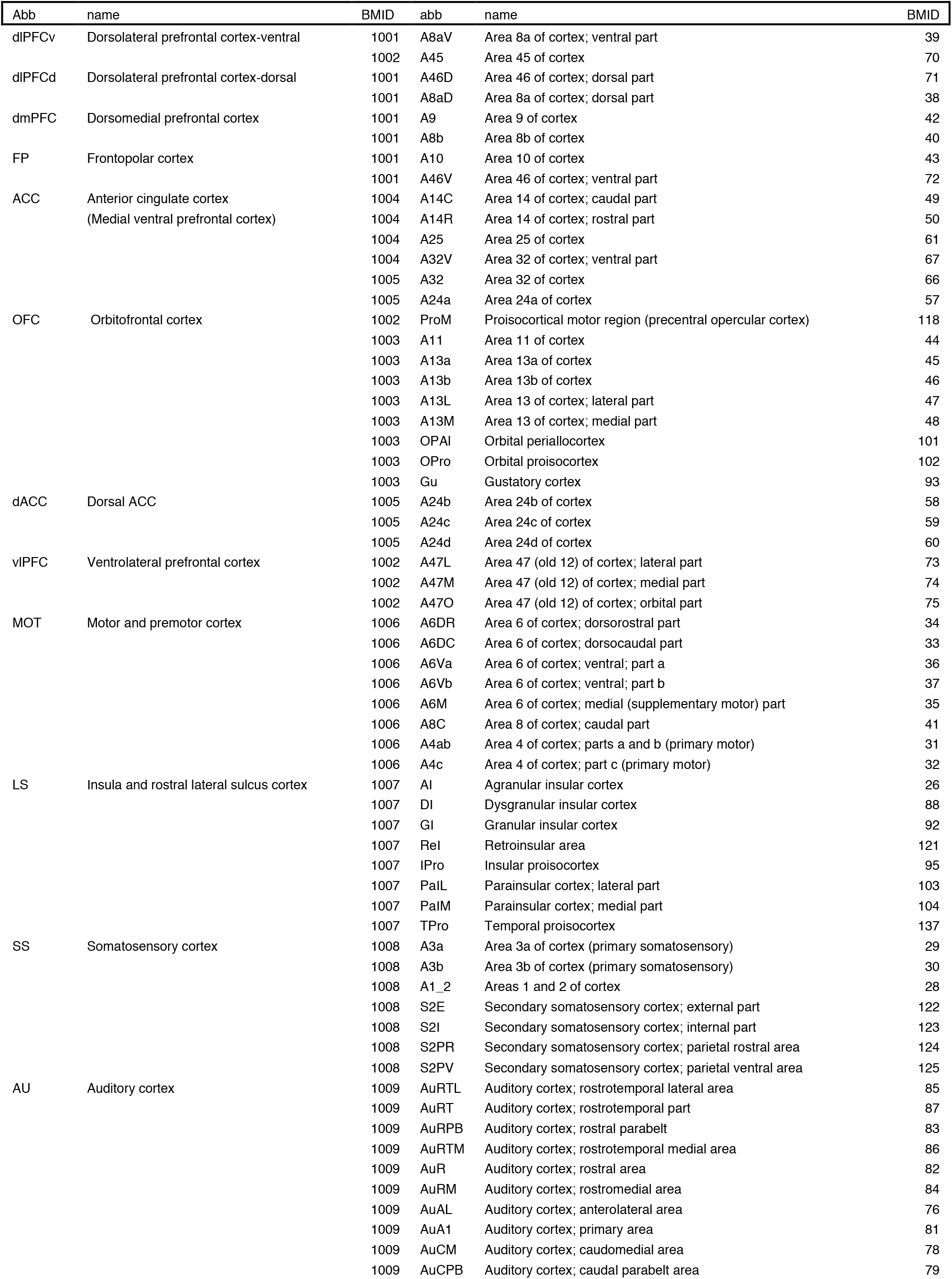
Marmoset cortical areas classified into area blocks.

**Supplementary Table2-2.**
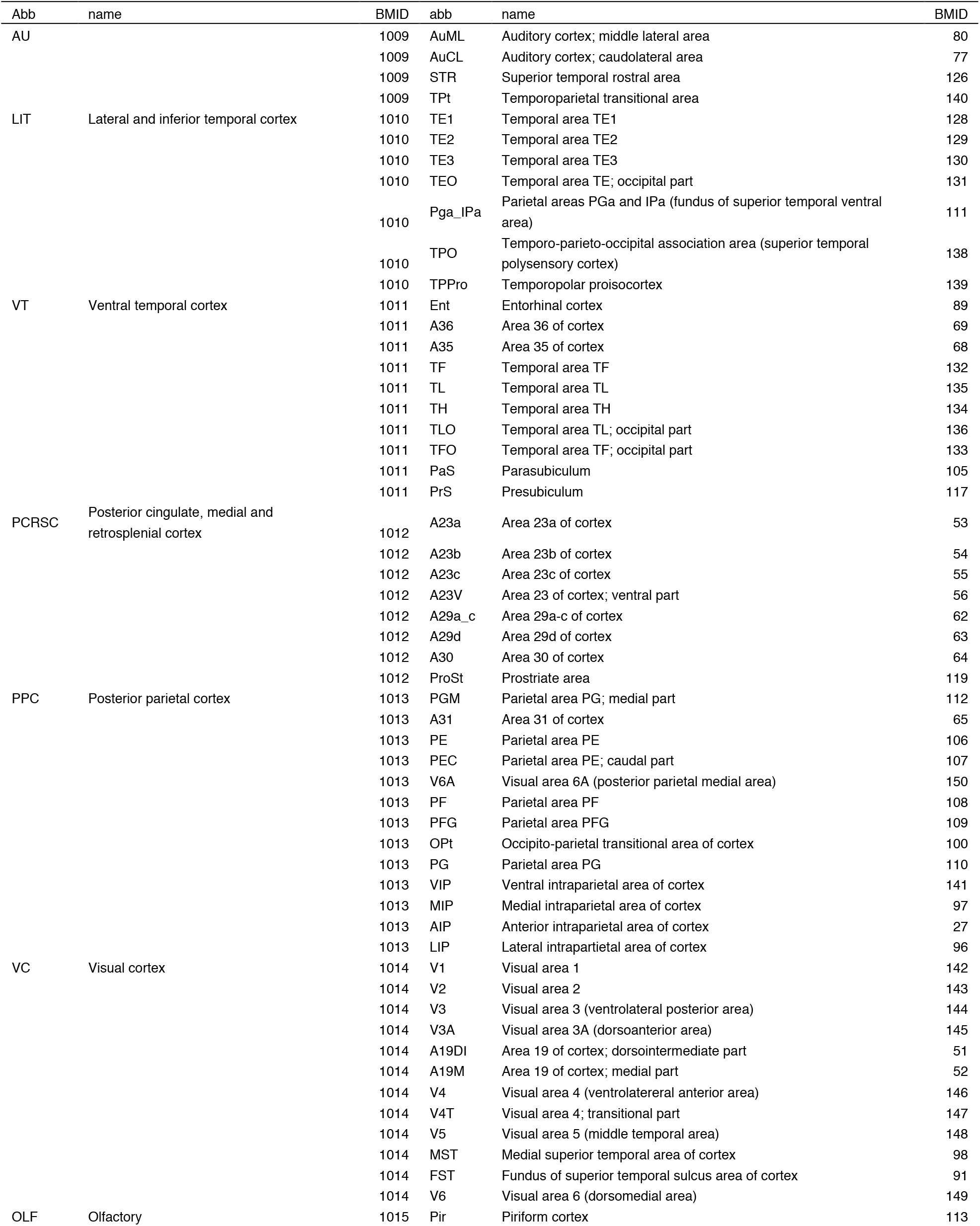
Marmoset cortical areas classified into area blocks. BMID: Brain/MINDS structural ID

**Supplementary Table3.** Mouse data analyzed in the present study.

| Data ID | injection | experiment_page_url |
| --- | --- | --- |
| 112458114 | ACAd | http://connectivity.brain-map.org/projection/experiment/112458114 |
| 139426984 | ACAd | http://connectivity.brain-map.org/projection/experiment/139426984 |
| 146593590 | ACAd | http://connectivity.brain-map.org/projection/experiment/146593590 |
| 112514202 | ACAv | http://connectivity.brain-map.org/projection/experiment/112514202 |
| 139520203 | ACAv | http://connectivity.brain-map.org/projection/experiment/139520203 |
| 100140756 | FRP | http://connectivity.brain-map.org/projection/experiment/100140756 |
| 157556400 | ILA | http://connectivity.brain-map.org/projection/experiment/157556400 |
| 100141454 | MOs | http://connectivity.brain-map.org/projection/experiment/100141454 |
| 112952510 | MOs | http://connectivity.brain-map.org/projection/experiment/112952510 |
| 141602484 | MOs | http://connectivity.brain-map.org/projection/experiment/141602484 |
| 141603190 | MOs | http://connectivity.brain-map.org/projection/experiment/141603190 |
| 157710335 | MOs | http://connectivity.brain-map.org/projection/experiment/157710335 |
| 180709942 | MOs | http://connectivity.brain-map.org/projection/experiment/180709942 |
| 180719293 | MOs | http://connectivity.brain-map.org/projection/experiment/180719293 |
| 180916954 | MOs | http://connectivity.brain-map.org/projection/experiment/180916954 |
| 585025284 | MOs | http://connectivity.brain-map.org/projection/experiment/585025284 |
| 112306316 | ORBI | http://connectivity.brain-map.org/projection/experiment/112306316 |
| 170721670 | ORBI | http://connectivity.brain-map.org/projection/experiment/170721670 |
| 180673746 | ORBI | http://connectivity.brain-map.org/projection/experiment/180673746 |
| 180709230 | ORBI | http://connectivity.brain-map.org/projection/experiment/180709230 |
| 126860974 | ORBm | http://connectivity.brain-map.org/projection/experiment/126860974 |
| 112423392 | ORBvl | http://connectivity.brain-map.org/projection/experiment/112423392 |
| 158435116 | ORBvl | http://connectivity.brain-map.org/projection/experiment/158435116 |
| 157711748 | PL | http://connectivity.brain-map.org/projection/experiment/157711748 |

## References

1. Fuster, J.M. (2015). The prefrontal cortex Fifth edition. (Elsevier/AP, Academic Press is an imprint of Elsevier).

2. Haber, S.N., Liu, H., Seidlitz, J., and Bullmore, E. (2021). Prefrontal connectomics: from anatomy to human imaging. Neuropsychopharmacology. 10.1038/s41386-021-01156-6.

3. Donahue, C.J., Glasser, M.F., Preuss, T.M., Rilling, J.K., and Van Essen, D.C. (2018). Quantitative assessment of prefrontal cortex in humans relative to nonhuman primates. Proc Natl Acad Sci U S A 115, E5183–E5192. 10.1073/pnas.1721653115.

4. Paxinos, G., Watson, C., Petrides, M., Rosa, M., and Tokuno, H. (2012). The marmoset brain in stereotaxic coordinates (Academic Press).

5. Majka, P., Bai, S., Bakola, S., Bednarek, S., Chan, J.M., Jermakow, N., Passarelli, L., Reser, D.H., Theodoni, P., Worthy, K.H., et al. (2020). Open access resource for cellular-resolution analyses of corticocortical connectivity in the marmoset monkey. Nat Commun 11, 1133. 10.1038/s41467-020-14858-0.

6. Pandya, D.N., Seltzer, B., Petrides, M., and Cipolloni, P.B. (2014). Cerebral cortex: architecture, connections, and the dual origin concept (Oxford University Press).

7. Markov, N.T., Vezoli, J., Chameau, P., Falchier, A., Quilodran, R., Huissoud, C., Lamy, C., Misery, P., Giroud, P., Ullman, S., et al. (2014). Anatomy of hierarchy: feedforward and feedback pathways in macaque visual cortex. J Comp Neurol 522, 225–259. 10.1002/cne.23458.

8. Vanni, S., Hokkanen, H., Werner, F., and Angelucci, A. (2020). Anatomy and Physiology of Macaque Visual Cortical Areas V1, V2, and V5/MT: Bases for Biologically Realistic Models. Cereb Cortex 30, 3483–3517. 10.1093/cercor/bhz322.

9. Horton, J.C., and Adams, D.L. (2005). The cortical column: a structure without a function. Philos Trans R Soc Lond B Biol Sci 360, 837–862. 10.1098/rstb.2005.1623.

10. Roe, A.W. (2019). Columnar connectome: toward a mathematics of brain function. Netw Neurosci 3, 779–791. 10.1162/netn_a_00088.

11. Bugbee, N.M., and Goldman-Rakic, P.S. (1983). Columnar organization of corticocortical projections in squirrel and rhesus monkeys: similarity of column width in species differing in cortical volume. J Comp Neurol 220, 355–364. 10.1002/cne.902200309.

12. Levitt, J.B., Lewis, D.A., Yoshioka, T., and Lund, J.S. (1993). Topography of pyramidal neuron intrinsic connections in macaque monkey prefrontal cortex (areas 9 and 46). J Comp Neurol 338, 360–376. 10.1002/cne.903380304.

13. Lund, J.S., Yoshioka, T., and Levitt, J.B. (1993). Comparison of intrinsic connectivity in different areas of macaque monkey cerebral cortex. Cereb Cortex 3, 148–162. 10.1093/cercor/3.2.148.

14. Pucak, M.L., Levitt, J.B., Lund, J.S., and Lewis, D.A. (1996). Patterns of intrinsic and associational circuitry in monkey prefrontal cortex. J Comp Neurol 376, 614–630. 10.1002/(SICI)1096-9861(19961223)376:4<614::AID-CNE9>3.0.CO;2-4.

15. Hori, Y., Cléry, J.C., Selvanayagam, J., Schaeffer, D.J., Johnston, K.D., Menon, R.S., and Everling, S. (2021). Interspecies activation correlations reveal functional correspondences between marmoset and human brain areas. Proc Natl Acad Sci U S A 118, e2110980118. 10.1073/pnas.2110980118.

16. Kaneko, T., Komatsu, M., Yamamori, T., Ichinohe, N., and Okano, H. (2022). Cortical neural dynamics unveil the rhythm of natural visual behavior in marmosets. Commun Biol 5, 108. 10.1038/s42003-022-03052-1.

17. Lin, M.K., Takahashi, Y.S., Huo, B.-X., Hanada, M., Nagashima, J., Hata, J., Tolpygo, A.S., Ram, K., Lee, B.C., Miller, M.I., et al. (2019). A high-throughput neurohistological pipeline for brain-wide mesoscale connectivity mapping of the common marmoset. Elife 8, e40042. 10.7554/eLife.40042.

18. Liu, C., Yen, C.C.-C., Szczupak, D., Ye, F.Q., Leopold, D.A., and Silva, A.C. (2019). Anatomical and functional investigation of the marmoset default mode network. Nat Commun 10, 1975. 10.1038/s41467-019-09813-7.

19. Liu, C., Ye, F.Q., Newman, J.D., Szczupak, D., Tian, X., Yen, C.C.-C., Majka, P., Glen, D., Rosa, M.G.P., Leopold, D.A., et al. (2020). A resource for the detailed 3D mapping of white matter pathways in the marmoset brain. Nat Neurosci 23, 271–280. 10.1038/s41593-019-0575-0.

20. Mitchell, J.F., and Leopold, D.A. (2015). The marmoset monkey as a model for visual neuroscience. Neurosci Res 93, 20–46. 10.1016/j.neures.2015.01.008.

21. Okano, H. (2021). Current Status of and Perspectives on the Application of Marmosets in Neurobiology. Annu Rev Neurosci 44, 27–48. 10.1146/annurev-neuro-030520-101844.

22. Woodward, A., Hashikawa, T., Maeda, M., Kaneko, T., Hikishima, K., Iriki, A., Okano, H., and Yamaguchi, Y. (2018). The Brain/MINDS 3D digital marmoset brain atlas. Sci Data 5, 180009. 10.1038/sdata.2018.9.

23. Xu, R., Bichot, N.P., Takahashi, A., and Desimone, R. (2022). The cortical connectome of primate lateral prefrontal cortex. Neuron 110, 312-327.e7. 10.1016/j.neuron.2021.10.018.

24. Skibbe, H., Rachmadi, M.F., Nakae, K., Gutierrez, C.E., Hata, J., Tsukada, H., Poon, C., Doya, K., Majka, P., Rosa, M.G.P., et al. (2022). The Brain/MINDS Marmoset Connectivity Atlas: exploring bidirectional tracing and tractography in the same stereotaxic space (Neuroscience) 10.1101/2022.06.06.494999.

25. PRIMatE Data and Resource Exchange (PRIME-DRE) Global Collaboration Workshop and Consortium. Electronic address: and PRIMatE Data and Resource Exchange (PRIME-DRE) Global Collaboration Workshop and Consortium (2022). Toward next-generation primate neuroscience: A collaboration-based strategic plan for integrative neuroimaging. Neuron 110, 16–20. 10.1016/j.neuron.2021.10.015.

26. Harris, J.A., Mihalas, S., Hirokawa, K.E., Whitesell, J.D., Choi, H., Bernard, A., Bohn, P., Caldejon, S., Casal, L., Cho, A., et al. (2019). Hierarchical organization of cortical and thalamic connectivity. Nature 575, 195–202. 10.1038/s41586-019-1716-z.

27. Oh, S.W., Harris, J.A., Ng, L., Winslow, B., Cain, N., Mihalas, S., Wang, Q., Lau, C., Kuan, L., Henry, A.M., et al. (2014). A mesoscale connectome of the mouse brain. Nature 508, 207–214. 10.1038/nature13186.

28. Ragan, T., Kadiri, L.R., Venkataraju, K.U., Bahlmann, K., Sutin, J., Taranda, J., Arganda-Carreras, I., Kim, Y., Seung, H.S., and Osten, P. (2012). Serial two-photon tomography for automated ex vivo mouse brain imaging. Nat Methods 9, 255–258. 10.1038/nmeth.1854.

29. Isseroff, A., Schwartz, M.L., Dekker, J.J., and Goldman-Rakic, P.S. (1984). Columnar organization of callosal and associational projections from rat frontal cortex. Brain Res 293, 213–223. 10.1016/0006-8993(84)91228-9.

30. Buckner, R.L., and Margulies, D.S. (2019). Macroscale cortical organization and a default-like apex transmodal network in the marmoset monkey. Nat Commun 10, 1976. 10.1038/s41467-019-09812-8.

31. Barbas, H. (2015). General cortical and special prefrontal connections: principles from structure to function. Annu Rev Neurosci 38, 269–289. 10.1146/annurev-neuro-071714-033936.

32. Felleman, D.J., and Van Essen, D.C. (1991). Distributed hierarchical processing in the primate cerebral cortex. Cereb Cortex 1, 1–47. 10.1093/cercor/1.1.1-a.

33. Rockland, K.S., and Pandya, D.N. (1979). Laminar origins and terminations of cortical connections of the occipital lobe in the rhesus monkey. Brain Res 179, 3–20. 10.1016/0006-8993(79)90485-2.

34. Vezoli, J., Vinck, M., Bosman, C.A., Bastos, A.M., Lewis, C.M., Kennedy, H., and Fries, P. (2021). Brain rhythms define distinct interaction networks with differential dependence on anatomy. Neuron, S0896-6273(21)00725-X. 10.1016/j.neuron.2021.09.052.

35. Foster, N.N., Barry, J., Korobkova, L., Garcia, L., Gao, L., Becerra, M., Sherafat, Y., Peng, B., Li, X., Choi, J.-H., et al. (2021). The mouse cortico-basal ganglia-thalamic network. Nature 598, 188–194. 10.1038/s41586-021-03993-3.

36. Haber, S.N. (2016). Corticostriatal circuitry. Dialogues Clin Neurosci 18, 7–21.

37. Haber, S.N., Kim, K.-S., Mailly, P., and Calzavara, R. (2006). Reward-related cortical inputs define a large striatal region in primates that interface with associative cortical connections, providing a substrate for incentive-based learning. J Neurosci 26, 8368–8376. 10.1523/JNEUROSCI.0271-06.2006.

38. Selemon, L.D., and Goldman-Rakic, P.S. (1985). Longitudinal Topography and lnterdigitation of Corticostriatal Projections in the Rhesus Monkey’. The Journal of Neuroscience 5, 19.

39. Averbeck, B.B., Lehman, J., Jacobson, M., and Haber, S.N. (2014). Estimates of projection overlap and zones of convergence within frontal-striatal circuits. J Neurosci 34, 9497–9505. 10.1523/JNEUROSCI.5806-12.2014.

40. Borra, E., Ferroni, C.G., Gerbella, M., Giorgetti, V., Mangiaracina, C., Rozzi, S., and Luppino, G. (2019). Rostro-caudal Connectional Heterogeneity of the Dorsal Part of the Macaque Prefrontal Area 46. Cereb Cortex 29, 485–504. 10.1093/cercor/bhx332.

41. Ferry, A.T., Ongür, D., An, X., and Price, J.L. (2000). Prefrontal cortical projections to the striatum in macaque monkeys: evidence for an organization related to prefrontal networks. J Comp Neurol 425, 447–470. 10.1002/1096-9861(20000925)425:3<447::aid-cne9>3.0.co;2-v.

42. Goldman-Rakic, P.S. (1988). Topography of cognition: parallel distributed networks in primate association cortex. Annu Rev Neurosci 11, 137–156. 10.1146/annurev.ne.11.030188.001033.

43. Roberts, A.C., Tomic, D.L., Parkinson, C.H., Roeling, T.A., Cutter, D.J., Robbins, T.W., and Everitt, B.J. (2007). Forebrain connectivity of the prefrontal cortex in the marmoset monkey (Callithrix jacchus): an anterograde and retrograde tract-tracing study. J Comp Neurol 502, 86–112. 10.1002/cne.21300.

44. Badre, D., and D’Esposito, M. (2009). Is the rostro-caudal axis of the frontal lobe hierarchical? Nat Rev Neurosci 10, 659–669. 10.1038/nrn2667.

45. Riley, M.R., Qi, X.-L., Zhou, X., and Constantinidis, C. (2018). Anterior-posterior gradient of plasticity in primate prefrontal cortex. Nat Commun 9, 3790. 10.1038/s41467-018-06226-w.

46. Huntenburg, J.M., Bazin, P.-L., and Margulies, D.S. (2018). Large-Scale Gradients in Human Cortical Organization. Trends Cogn Sci 22, 21–31. 10.1016/j.tics.2017.11.002.

47. Du, J., and Buckner, R.L. (2021). Precision estimates of macroscale network organization in the human and their relation to anatomical connectivity in the marmoset monkey. Current Opinion in Behavioral Sciences 40, 144–152. 10.1016/j.cobeha.2021.04.010.

48. Braga, R.M., and Buckner, R.L. (2017). Parallel Interdigitated Distributed Networks within the Individual Estimated by Intrinsic Functional Connectivity. Neuron 95, 457-471.e5. 10.1016/j.neuron.2017.06.038.

49. Luo, L., Callaway, E.M., and Svoboda, K. (2018). Genetic Dissection of Neural Circuits: A Decade of Progress. Neuron 98, 256–281. 10.1016/j.neuron.2018.03.040.

50. Xu, X., Holmes, T.C., Luo, M.-H., Beier, K.T., Horwitz, G.D., Zhao, F., Zeng, W., Hui, M., Semler, B.L., and Sandri-Goldin, R.M. (2020). Viral Vectors for Neural Circuit Mapping and Recent Advances in Trans-synaptic Anterograde Tracers. Neuron 107, 1029–1047. 10.1016/j.neuron.2020.07.010.

51. Li, X., Zhu, Q., and Vanduffel, W. (2022). Submillimeter fMRI reveals an extensive, fine-grained and functionally-relevant scene-processing network in monkeys. Prog Neurobiol 211, 102230. 10.1016/j.pneurobio.2022.102230.

52. LeVay, S., Connolly, M., Houde, J., and Van Essen, D.C. (1985). The complete pattern of ocular dominance stripes in the striate cortex and visual field of the macaque monkey. J Neurosci 5, 486–501.

53. Lein, E., Borm, L.E., and Linnarsson, S. (2017). The promise of spatial transcriptomics for neuroscience in the era of molecular cell typing. Science 358, 64–69. 10.1126/science.aan6827.

54. Sincich, L.C., and Horton, J.C. (2005). The circuitry of V1 and V2: integration of color, form, and motion. Annu Rev Neurosci 28, 303–326. 10.1146/annurev.neuro.28.061604.135731.

55. Bao, P., She, L., McGill, M., and Tsao, D.Y. (2020). A map of object space in primate inferotemporal cortex. Nature 583, 103–108. 10.1038/s41586-020-2350-5.

56. Fujita, I., Tanaka, K., Ito, M., and Cheng, K. (1992). Columns for visual features of objects in monkey inferotemporal cortex. Nature 360, 343–346. 10.1038/360343a0.

57. Liu, Y., Li, M., Zhang, X., Lu, Y., Gong, H., Yin, J., Chen, Z., Qian, L., Yang, Y., Andolina, I.M., et al. (2020). Hierarchical Representation for Chromatic Processing across Macaque V1, V2, and V4. Neuron 108, 538-550.e5. 10.1016/j.neuron.2020.07.037.

58. Mante, V., Sussillo, D., Shenoy, K.V., and Newsome, W.T. (2013). Context-dependent computation by recurrent dynamics in prefrontal cortex. Nature 503, 78–84. 10.1038/nature12742.

59. Rigotti, M., Barak, O., Warden, M.R., Wang, X.-J., Daw, N.D., Miller, E.K., and Fusi, S. (2013). The importance of mixed selectivity in complex cognitive tasks. Nature 497, 585–590. 10.1038/nature12160.

60. Stringer, C., Pachitariu, M., Steinmetz, N., Reddy, C.B., Carandini, M., and Harris, K.D. (2019). Spontaneous behaviors drive multidimensional, brainwide activity. Science 364, 255. 10.1126/science.aav7893.

61. Watakabe, A., Ohtsuka, M., Kinoshita, M., Takaji, M., Isa, K., Mizukami, H., Ozawa, K., Isa, T., and Yamamori, T. (2015). Comparative analyses of adeno-associated viral vector serotypes 1, 2, 5, 8 and 9 in marmoset, mouse and macaque cerebral cortex. Neurosci Res 93, 144–157. 10.1016/j.neures.2014.09.002.

62. Hioki, H., Kuramoto, E., Konno, M., Kameda, H., Takahashi, Y., Nakano, T., Nakamura, K.C., and Kaneko, T. (2009). High-level transgene expression in neurons by lentivirus with Tet-Off system. Neurosci Res 63, 149–154. 10.1016/j.neures.2008.10.010.

63. Sadakane, O., Watakabe, A., Ohtsuka, M., Takaji, M., Sasaki, T., Kasai, M., Isa, T., Kato, G., Nabekura, J., Mizukami, H., et al. (2015). In Vivo Two-Photon Imaging of Dendritic Spines in Marmoset Neocortex. eNeuro 2, ENEURO.0019-15.2015. 10.1523/ENEURO.0019-15.2015.

64. Sadakane, O., Masamizu, Y., Watakabe, A., Terada, S.-I., Ohtsuka, M., Takaji, M., Mizukami, H., Ozawa, K., Kawasaki, H., Matsuzaki, M., et al. (2015). Long-Term Two-Photon Calcium Imaging of Neuronal Populations with Subcellular Resolution in Adult Non-human Primates. Cell Rep 13, 1989–1999. 10.1016/j.celrep.2015.10.050.

65. Watakabe, A., Takaji, M., Kato, S., Kobayashi, K., Mizukami, H., Ozawa, K., Ohsawa, S., Matsui, R., Watanabe, D., and Yamamori, T. (2014). Simultaneous visualization of extrinsic and intrinsic axon collaterals in Golgi-like detail for mouse corticothalamic and corticocortical cells: a double viral infection method. Front Neural Circuits 8, 110. 10.3389/fncir.2014.00110.

66. Reser, D.H., Burman, K.J., Yu, H.-H., Chaplin, T.A., Richardson, K.E., Worthy, K.H., and Rosa, M.G.P. (2013). Contrasting patterns of cortical input to architectural subdivisions of the area 8 complex: a retrograde tracing study in marmoset monkeys. Cereb Cortex 23, 1901–1922. 10.1093/cercor/bhs177.

67. Burman, K.J., Reser, D.H., Yu, H.-H., and Rosa, M.G.P. (2011). Cortical input to the frontal pole of the marmoset monkey. Cereb Cortex 21, 1712–1737. 10.1093/cercor/bhq239.

68. Alexander, L., Gaskin, P.L.R., Sawiak, S.J., Fryer, T.D., Hong, Y.T., Cockcroft, G.J., Clarke, H.F., and Roberts, A.C. (2019). Fractionating Blunted Reward Processing Characteristic of Anhedonia by Over-Activating Primate Subgenual Anterior Cingulate Cortex. Neuron 101, 307-320.e6. 10.1016/j.neuron.2018.11.021.

69. Selvanayagam, J., Johnston, K.D., Schaeffer, D.J., Hayrynen, L.K., and Everling, S. (2019). Functional Localization of the Frontal Eye Fields in the Common Marmoset Using Microstimulation. J Neurosci 39, 9197–9206. 10.1523/JNEUROSCI.1786-19.2019.

70. Eldred, G.E., Miller, G.V., Stark, W.S., and Feeney-Burns, L. (1982). Lipofuscin: resolution of discrepant fluorescence data. Science 216, 757–759. 10.1126/science.7079738.

71. Skibbe, H., Watakabe, A., Nakae, K., Gutierrez, C.E., Tsukada, H., Hata, J., Kawase, T., Gong, R., Woodward, A., Doya, K., et al. (2019). MarmoNet: a pipeline for automated projection mapping of the common marmoset brain from whole-brain serial two-photon tomography. arXiv:1908.00876 [cs, eess, q-bio, stat].

72. Woodward, A., Gong, R., Abe, H., Nakae, K., Hata, J., Skibbe, H., Yamaguchi, Y., Ishii, S., Okano, H., Yamamori, T., et al. (2020). The NanoZoomer artificial intelligence connectomics pipeline for tracer injection studies of the marmoset brain. Brain Struct Funct 225, 1225–1243. 10.1007/s00429-020-02073-y.

73. Iriki, A., Okano, J. H., Sasaki, E., and Okano, H. (2018). The 3-dimensional atlas of the marmoset brain (Springer Berlin Heidelberg).

74. Fedorov, A., Beichel, R., Kalpathy-Cramer, J., Finet, J., Fillion-Robin, J.-C., Pujol, S., Bauer, C., Jennings, D., Fennessy, F., Sonka, M., et al. (2012). 3D Slicer as an image computing platform for the Quantitative Imaging Network. Magn Reson Imaging 30, 1323–1341. 10.1016/j.mri.2012.05.001.

75. Wan, Y., Otsuna, H., Holman, H.A., Bagley, B., Ito, M., Lewis, A.K., Colasanto, M., Kardon, G., Ito, K., and Hansen, C. (2017). FluoRender: joint freehand segmentation and visualization for many-channel fluorescence data analysis. BMC Bioinformatics 18, 280. 10.1186/s12859-017-1694-9.

76. Avants, B.B., Tustison, N.J., Song, G., Cook, P.A., Klein, A., and Gee, J.C. (2011). A reproducible evaluation of ANTs similarity metric performance in brain image registration. Neuroimage 54, 2033–2044. 10.1016/j.neuroimage.2010.09.025.

77. Natan (2021). Fast 2D peak finder (https://www.mathworks.com/matlabcentral/fileexchange/37388-fast-2d-peak-finder), MATLAB Central File Exchange. Retrieved May 26, 2021.

78. Weed, N., Bakken, T., Graddis, N., Gouwens, N., Millman, D., Hawrylycz, M., and Waters, J. (2019). Identification of genetic markers for cortical areas using a Random Forest classification routine and the Allen Mouse Brain Atlas. PLoS One 14, e0212898. 10.1371/journal.pone.0212898.

79. Sotiras, A., Resnick, S.M., and Davatzikos, C. (2015). Finding imaging patterns of structural covariance via Non-Negative Matrix Factorization. Neuroimage 108, 1–16. 10.1016/j.neuroimage.2014.11.045.

80. Lee, D.D., and Seung, H.S. (1999). Learning the parts of objects by non-negative matrix factorization. Nature 401, 788–791. 10.1038/44565.

81. Steiger, M., Bernard, J., Mittelstädt, S., Hutter, M., Keim, D., Thum, S., and Kohlhammer, J. (2015). Explorative analysis of 2D color maps. In Proceedings of WSCG, pp. 151–160.

82. Schindelin, J., Arganda-Carreras, I., Frise, E., Kaynig, V., Longair, M., Pietzsch, T., Preibisch, S., Rueden, C., Saalfeld, S., Schmid, B., et al. (2012). Fiji: an open-source platform for biological-image analysis. Nat Methods 9, 676–682. 10.1038/nmeth.2019.

83. Rutledge, R.B., and Adams, R.A. (2017). Computational Psychiatry. In Computational Models of Brain and Behavior (John Wiley & Sons, Ltd.).

84. Greicius, M.D., Srivastava, G., Reiss, A.L., and Menon, V. (2004). Default-mode network activity distinguishes Alzheimer’s disease from healthy aging: evidence from functional MRI. Proc Natl Acad Sci U S A 101, 4637–4642. 10.1073/pnas.0308627101.

85. Li, T., Wang, Q., Zhang, J., Rolls, E.T., Yang, W., Palaniyappan, L., Zhang, L., Cheng, W., Yao, Y., Liu, Z., et al. (2017). Brain-Wide Analysis of Functional Connectivity in First-Episode and Chronic Stages of Schizophrenia. Schizophr Bull 43, 436–448. 10.1093/schbul/sbw099.

86. Northoff, G., and Duncan, N.W. (2016). How do abnormalities in the brain’s spontaneous activity translate into symptoms in schizophrenia? From an overview of resting state activity findings to a proposed spatiotemporal psychopathology. Prog Neurobiol 145–146, 26–45. 10.1016/j.pneurobio.2016.08.003.

